# Multimodal autonomic arousal tracks dose-dependent affective dynamics during the acute effects of DMT

**DOI:** 10.64898/2026.04.30.721872

**Authors:** Tomás Ariel D’Amelio, Tomás Gil Garbagnoli, Jerónimo Rodríguez Cuello, Evan Lewis-Healey, Carla Pallavicini, Federico Cavanna, Nicolás Marcelo Bruno, Laura Alethia De La Fuente, Stephanie Müller, Débora Copa, Tristan Bekinschtein, Diego Vidaurre, Enzo Tagliazucchi

## Abstract

Serotonergic psychedelics induce altered states of consciousness characterised by profound changes in emotional experience. Although psychedelics modulate autonomic arousal, sympathetic engagement during their affective effects remains poorly characterised. We recorded cardiac, electrodermal, and respiratory activity in 19 participants inhaling 20 or 40 mg of N,N-dimethyltryptamine (DMT) under a semi-naturalistic blinded design, alongside time-resolved retrospective phenomenological reports. DMT increased heart rate and sudomotor activity, with a transient dose-dependent rise in respiratory volume. A multimodal index integrating the three signals selectively tracked subjective emotional intensity. Dose-dependent divergence followed modality-specific profiles: heart rate and respiratory differences emerged within the first 2 min post-inhalation, whereas electrodermal activity diverged only during the later phase. At 40 mg, peak heart rate exceeded the rate attainable by vagal withdrawal alone in most participants, indicating active sympathetic engagement. DMT thus produces a transient sympathetic activation covarying with emotional arousal, followed by gradual disengagement accompanied by pleasantness and bliss. By combining time- and cost-effective peripheral physiological measures with time-resolved phenomenological reports, this work contributes to the objective characterisation of psychedelic-induced affective states and provides a methodological basis for future biomarker research in clinical applications.

---

Serotonergic psychedelics act primarily as partial agonists at serotonin receptors, most notably the 5-HT_2A_ receptor, and are remarkable for their capacity to induce profound yet transient alterations in the contents of consciousness^1–3^. Their subjective effects unfold along multiple dimensions, including changes in perception^4^, self- and bodily awareness^5^, and the processing of emotions^6^. Among psychedelics, N,N-dimethyltryptamine (DMT) is distinguished by its rapid onset and short duration of action, making it particularly well suited for investigating the dynamic transitions between states of consciousness and their underlying neural correlates^7–10^. Growing evidence also supports the clinical potential of DMT and other psychedelics in the treatment of mood disorders^11–15^, underscoring the need to better understand how these substances influence affective processing, how such effects relate to activity in the central and autonomic nervous systems, and how transient psychedelic experiences may lead to lasting improvements in emotional well-being.

Emotions can be conceptualised as internal representations of salient extrinsic or intrinsic stimuli that guide behaviour^16^. Numerous theoretical models of emotion have been proposed, ranging from discrete categorical accounts to dimensional frameworks based on latent continuous variables^17^. One of the most influential dimensional frameworks is the Circumplex Model of Emotion^18^, which posits that affective states can be organised within a two-dimensional circular space defined by valence (pleasantness–unpleasantness) and arousal (low to high intensity). As early as 1884, William James emphasised the embodied nature of emotion, writing that “bodily changes follow directly the perception of the exciting fact, and that our feeling of the same changes as they occur is the emotion”^19^. Consistent with this view, substantial empirical evidence supports a close relationship between autonomic nervous system (ANS) activity and affective processing^20,21^. Physiological measures of cardiac and electrodermal activity have therefore been widely investigated to detect the circumplex dimensions of arousal and valence, reinforcing the notion that emotional experience is intrinsically linked to autonomic physiology^22–24^.

The ANS consists of two primary and interacting branches: the sympathetic nervous system, which supports physiological reactivity to salient or potentially threatening situations, and the parasympathetic nervous system, which counterbalances sympathetic activity and promotes homeostasis^25^. Subjective reports of psychedelic-induced affective states suggest a dynamic interplay between these two branches. Following the onset of acute effects, users often report an initial phase characterised by heightened arousal, restlessness, or anxiety, which can gradually subside into states of relaxation and enhanced emotional well-being^1,26^. This phenomenological time course has been interpreted as reflecting an initial engagement of sympathetic processes, potentially accompanied by parasympathetic withdrawal, followed by increased parasympathetic involvement during later stages of the experience^27^. Consistent with this view, previous studies have documented sympathetic effects of several psychedelics, including elevations in heart rate (HR), blood pressure, and pupil diameter^28–30^. A post-stress vagal rebound, which would appear as enhanced parasympathetic modulation of cardiac activity during later phases of the experience, has further been proposed to follow this initial sympathetic engagement^31^. Together, findings from DMT and other psychedelic studies point to a joint and time-varying influence of sympathetic and parasympathetic processes on cardiovascular dynamics, with early sympathetic predominance followed by a gradual shift towards parasympathetic regulation.

However, the relationship between psychedelic-induced changes in ANS activity and affective states remains poorly understood due to two key methodological limitations in prior research. First, previous studies have relied primarily on cardiac measures as indices of ANS activity; yet such measures are inherently underdetermined with respect to the selective engagement of sympathetic and parasympathetic branches^32^. Although heart rate variability (HRV) has frequently been interpreted as reflecting increased vagal tone, its estimation can be substantially confounded by the pronounced non-stationary cardiorespiratory dynamics induced by DMT^26,28,33,34^. Second, the correspondence between physiological signals and affective experience has been further limited by the lack of continuous subjective reports of emotional valence and arousal^35^. As a result, it remains unclear whether strong sympathetic activation is present at the onset of the DMT experience, and how any such activation, and its subsequent attenuation, relates to the evolving emotional states reported by participants across different phases of the experience.

In this study, we leveraged the short-lived effects of two doses of DMT (20 mg and 40 mg) to investigate the relationship between ANS activity and the temporal dynamics of affective states (Fig. 1). In addition to electrocardiogram (ECG) recordings of heart rate (HR), we incorporated complementary physiological measures, including sudomotor nerve activity (SMNA) indexed by electrodermal activity (EDA) and respiratory volume per time (RVT) measured using a respiratory belt. This multimodal approach enables a more comprehensive characterisation of autonomic arousal, with improved specificity for sympathetic activation. Heart rate can rise either through sympathetic drive or through the release of vagal restraint. To separate the two, we compared each participant’s peak heart rate with their estimated intrinsic rate, the rate of the heart without autonomic input, which vagal withdrawal alone cannot exceed^36–38^. To track the dynamic evolution of affective states, we employed the recently introduced Temporal Experience Tracing (TET) method. TET is a phenomenological approach in which participants retrospectively trace the subjective intensity of experiential dimensions over time, yielding continuous time-resolved data rather than discrete summary scores derived from standard psychometric questionnaires^10,39–44^. In the present study, we focused on a predefined subset of affective TET dimensions—Pleasantness, Unpleasantness, Emotional Intensity, Interoception, Bliss, and Anxiety—to characterise the temporal dynamics of emotional experience. Together, these measures allow us to ask whether the cardiovascular response to DMT reflects active sympathetic engagement rather than the release of parasympathetic restraint alone, how the time course of autonomic activity differs across effectors, and how that activity relates to the unfolding affective experience.

**Figure 1:**
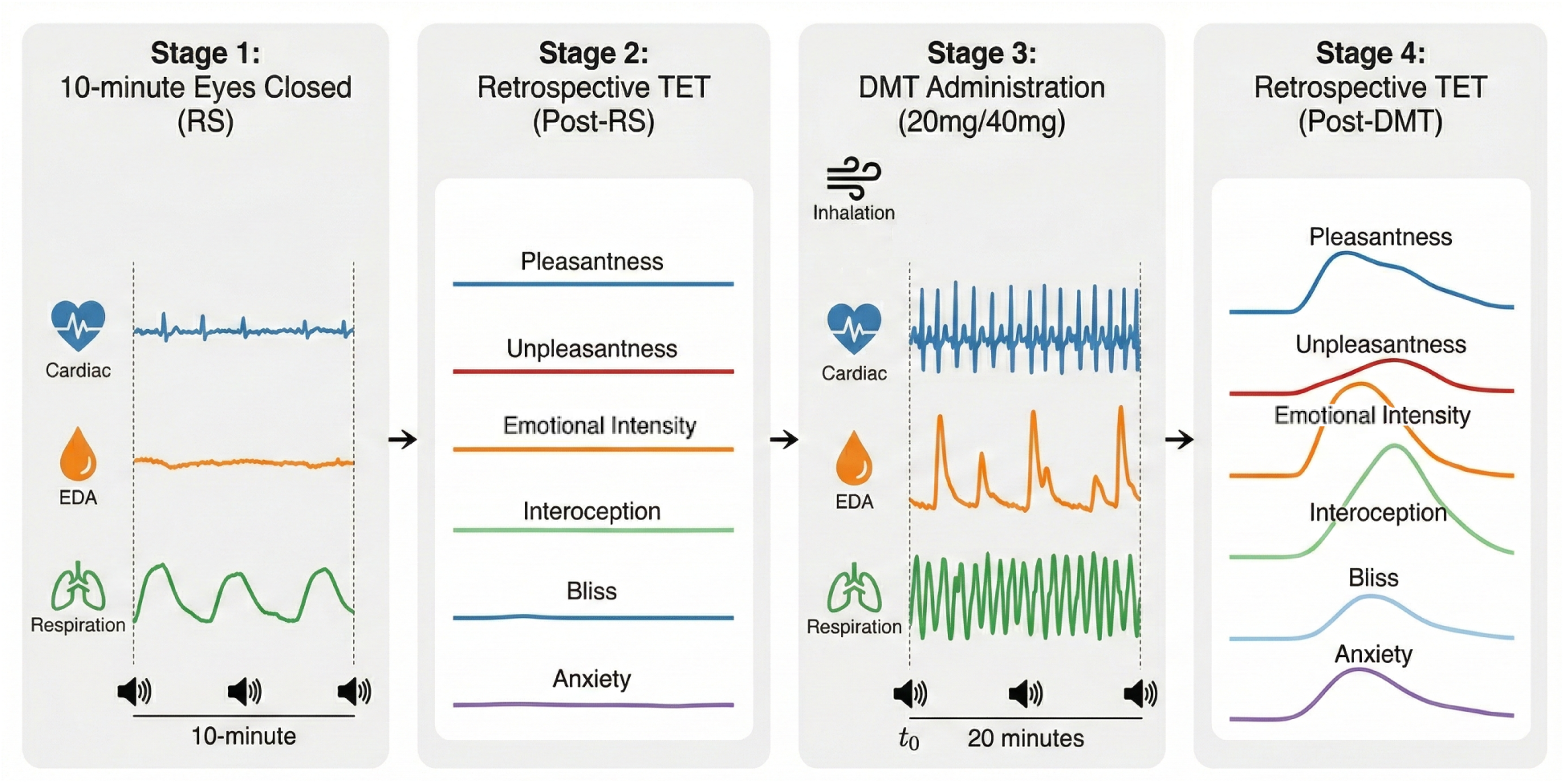
Experimental timeline. Each session comprised four stages: a 10-min eyes-closed Resting State (RS) baseline with continuous physiological recording (Stage 1), retrospective Temporal Experience Tracing (TET) of the resting-state experience (Stage 2), DMT inhalation (20 mg or 40 mg freebase) followed by 20 min of recording (Stage 3), and retrospective TET of the DMT experience (Stage 4). Auditory chimes every two minutes served as temporal anchors for retrospective tracing. Stages 2 and 4 display the six affective TET dimensions analysed in this study; participants completed the full 15-dimension battery (see Methods § Affective phenomenology). The recording setup and the signal processing pipeline are shown in Supplementary Fig. 1.

Based on studies of DMT and other serotonergic psychedelics, we formulated the following hypotheses: 1) Both moderate (20 mg) and high (40 mg) doses of freebase DMT induce robust changes in HR, SMNA, and RVT, characterised by a rapid increase at the onset of the experience consistent with sympathetic nervous system engagement, 2) A multimodal index of autonomic arousal that integrates these physiological measures exhibits greater sensitivity to the temporal dynamics and dose-dependent effects of DMT, 3) The latent structure of a predefined subset of affective TET dimensions (Pleasantness, Unpleasantness, Emotional Intensity, Interoception, Bliss, and Anxiety) reveals components corresponding to emotional arousal and valence, with an initial phase of heightened arousal followed by a reduction in arousal and a shift towards more positive valence over time, and 4) The multimodal index of physiological arousal is positively associated with subjective reports of arousal during the DMT experience^7,9,10,31,33,45^.

## Results

We first examined the dose-dependent effects of DMT on individual autonomic measures (heart rate, electrodermal activity, and respiration) and their shared latent structure, followed by the characterisation of affective dynamics from time-resolved subjective reports. We then assessed the coupling between physiological activation and subjective experience using correlational and multivariate approaches. Individual-participant trajectories for heart rate, sudomotor nerve activity, and respiratory volume per time are shown in Supplementary Fig. 2.

### Electrocardiography

To assess cardiac responses to DMT, we examined heart rate (HR) across doses and states. Linear mixed-effects (LME) modelling revealed a robust State × Dose interaction, *β* = 1.12, 95% CI [0.87, 1.38], *p*_FDR_ *<* .001 (Fig. 2b), driven by a specific increase in HR during the DMT state (State effect: *β* = 0.69, 95% CI [0.46, 0.91], *p*_FDR_ *<* .001) that was attenuated in the Low dose condition. Every participant showed the interaction in the same direction (11 of 11; *M* = 1.12, *SD* = 0.37, Cohen’s *d* = 3.01). Conditional contrasts showed a large dose effect specific to the DMT state (High vs. Low: *β* = 0.85, Cohen’s *d* = 1.25). Throughout, the Dose coefficient of the LME models reflects the High vs. Low difference during the Resting State, which is the reference level, whereas the net dose effect during DMT is given by the sum of the Dose and State × Dose coefficients. For HR this coefficient was negative (*β* = − 0.27, *p*_FDR_ = .043; Fig. 2b). Because both baselines were recorded before inhalation, this difference cannot reflect the dose; it formed no cluster over time (two-tailed; Fig. 2a) and was not apparent in absolute resting heart rate (see below). In contrast, during the DMT state, a single sustained cluster of High *>* Low separation emerged approximately 1.5 min after inhalation and persisted throughout the 9-min main analysis window (cluster mass = 55.6, exact *p* = .0015; Fig. 2a), with the High dose tracking consistently above the Low dose; the effect gradually attenuated over the subsequent ten minutes (Supplementary Fig. 3).

**Figure 2:**
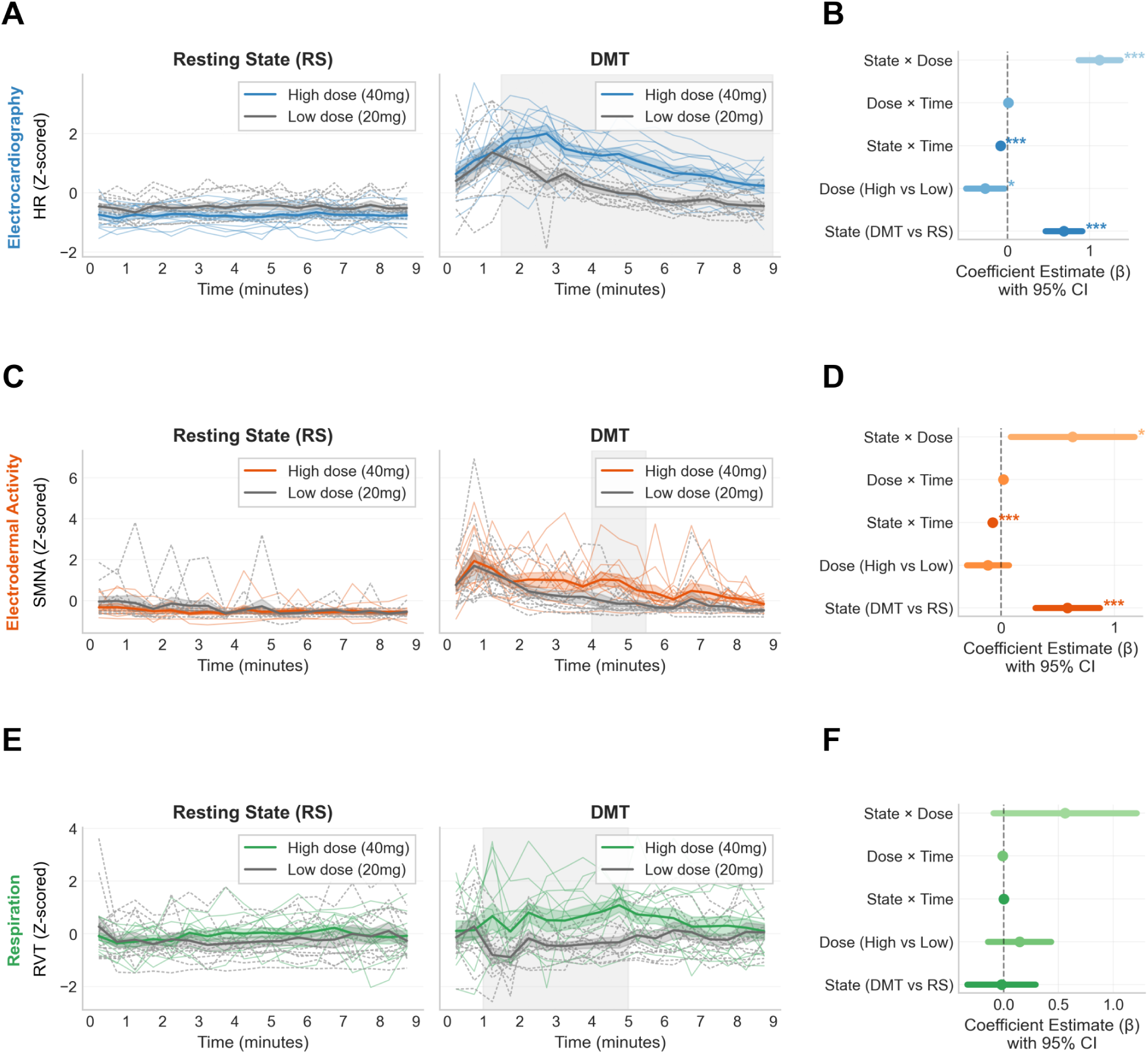
Dose-dependent autonomic activation following DMT inhalation. **a**, Heart rate (HR; *z*-scored) time courses during Resting State (RS) and DMT (mean ± s.e.m.; *n* = 11). Data are aligned to inhalation onset (*t*_0_). In **a**, **c**, and **e**, thin lines show each participant (High dose in colour, Low dose in grey, dashed) and shaded rectangles indicate clusters of significant High vs. Low separation (cluster-based sign-flip permutation tests; one-tailed for DMT, two-tailed for RS; see Methods). **b**, Fixed-effect coefficients (*β* and 95% CI) from the Linear Mixed-Effects (LME) model for HR (30-s bins; see Methods). Asterisks indicate significant terms (* *p <* .05, *** *p <* .001; FDR-corrected). **c**, Sudomotor nerve activity (SMNA; *z*-scored; *n* = 11) time courses. SMNA was computed as the area under the curve (AUC) within 30-s bins. **d**, LME coefficients for SMNA. **e**, Respiratory volume per time (RVT; *z*-scored; *n* = 12) time courses. **f**, LME coefficients for RVT. Source data are provided as a Source Data file (Supplementary Data 1).

Because the modelling above is carried out on within-participant *z*-scores, we also report heart rate in absolute units. Averaged over the 9-min window, resting heart rate was *M* = 65.8 bpm (*SD* = 9.4) before the High dose and *M* = 68.0 bpm (*SD* = 9.6) before the Low dose, a difference that was not significant (*p* = .436). Peak heart rate, taken over 60-s windows, reached *M* = 113.6 bpm (*SD* = 22.6; range 73.7–154.6) under the High dose and *M* = 99.2 bpm (*SD* = 12.9; range 86.4–125.2) under the Low dose, *t*(10) = 2.76, *p* = .020. Expressed as a rise above each participant’s own resting level, the High dose produced an increase of *M* = 47.8 bpm (*SD* = 18.5) against *M* = 31.2 bpm (*SD* = 14.7) for the Low dose, *t*(10) = 3.92, *p* = .003.

Each participant’s peak heart rate was compared with their intrinsic rate, estimated from age (Methods). The estimated intrinsic rate averaged 98.2 bpm (*SD* = 2.4; age range 27–42 years), well above the observed resting rates. Under the High dose, 8 of 11 participants exceeded their own estimated intrinsic rate, and 4 of 11 exceeded even the upper 95% limit of that estimate. Across participants, peak heart rate exceeded the estimated intrinsic rate by a mean of 15.4 bpm (median 11.9 bpm; maximum 57.6 bpm; *t*(10) = 2.28, *p* = .023; Wilcoxon *p* = .021; one-tailed). Under the Low dose the same comparison yielded 5 of 11 participants and a mean margin of 1.0 bpm (median 0.2 bpm; *p* = .398). The three participants who did not exceed their intrinsic rate under the High dose were those with the smallest cardiac response overall.

Time-domain heart rate variability was examined directly and is reported in full in the Supplementary Information (Supplementary Fig. 4). Briefly, the root mean square of successive differences (RMSSD) and the standard deviation of inter-beat intervals (SDNN) both decreased under DMT in a dose-dependent manner, and RMSSD fell below its resting level between 1 and 3 min after inhalation; however, once RMSSD was expressed relative to the mean inter-beat interval the effect was no longer detectable, indicating that the decrease in absolute variability tracked the shortening of the cardiac cycle and cannot be taken as evidence of vagal withdrawal over and above the heart rate change itself.

### Electrodermal activity

We next examined sympathetic activation through electrodermal activity, quantified as sudomotor nerve activity (SMNA). SMNA exhibited a dose-dependent amplification relative to rest (Fig. 2c). This was substantiated by a State × Dose interaction, *β* = 0.63, 95% CI [0.09, 1.17], *p*_FDR_ = .023 (Fig. 2d). While the main effect of State indicated increased sympathetic activation during DMT (*β* = 0.59, 95% CI [0.30, 0.87], *p*_FDR_ *<* .001), the interaction suggests this effect was stronger in the High dose condition. The interval around the interaction is wide because the size of the electrodermal dose response varies markedly across participants: per-participant State × Dose contrasts ranged from 1.02 to 1.59 (*M* = 0.63, *SD* = 0.87, Cohen’s *d* = 0.73), with seven of eleven participants positive. Consistent with this, the cluster-based permutation test identified a cluster of dose separation at approximately 4–5.5 min post-inhalation (cluster mass = 7.1, exact *p* = .041; Fig. 2c, shaded region). A second epoch towards the end of the analysis window, at 8–9 min, showed the same direction but did not reach significance (cluster mass = 5.3, exact *p* = .058). Unlike the single sustained cluster observed for HR, which began 1.5 min after inhalation and ran to the end of the window, electrodermal dose separation was therefore confined to a 1.5-min epoch beginning 4 min after inhalation. Window-wise FDR-corrected comparisons are reported in Supplementary Table 1.

### Respiration

We then assessed whether DMT modulated respiratory drive, measured as respiratory volume per time (RVT). RVT separated the doses transiently rather than on average (Fig. 2e). Averaged over the 9-min window, the State × Dose interaction was not significant, *β* = 0.56, 95% CI [ −0.09, 1.21], *p*_FDR_ = .093 (Fig. 2f), nor was the main effect of Dose (*β* = 0.14, *p*_FDR_ = .381). The effect is nonetheless present and localised in time: the cluster-based permutation test identified a single sustained cluster of dose separation spanning approximately 1–5 min post-inhalation (cluster mass = 21.4, exact *p* = .011; Fig. 2e, shaded region), and ten of twelve participants showed a positive contrast (*M* = 0.56, *SD* = 1.01, Cohen’s *d* = 0.55). Respiratory dose separation is therefore supported as a transient effect confined to the first five minutes, but not as an average difference across the whole window; the discrepancy is likely to reflect the variability of the respiratory response across participants, one of whom showed a contrast of 1.99 against a group mean of 0.56. Window-wise FDR-corrected comparisons, which localise the effect to individual 30-s windows, are reported in Supplementary Table 1.

Likelihood-ratio tests confirmed that the heteroscedastic model provided a significantly better fit for all three modalities (HR: *χ*^2^(1) = 247.9; SMNA: *χ*^2^(1) = 128.8; RVT: *χ*^2^(1) = 20.7; all *p <* .001), with residual standard deviations during DMT approximately 2.3, 1.8, and 1.3 times larger than during RS for HR, SMNA, and RVT, respectively. Crucially, the State, Dose, State × Dose, and State × Time coefficients were unchanged to three decimal places across standard and heteroscedastic models in all modalities; the Time and Dose × Time terms shifted slightly, retaining their sign and significance. The reported findings are therefore robust to this assumption.

### Latent component of autonomic arousal

To determine whether the three autonomic signals reflected a common underlying activation, we combined them into a single index using principal component analysis. The first principal component (PC1) of the joint physiological space (HR, SMNA, RVT), computed in the subset of participants with complete concurrent data (*n* = 7), was retained as the Physiological Arousal Index. This Physiological Arousal Index explained 58.9% of total variance and loaded positively on all modalities (loadings: HR = 0.63, SMNA = 0.64, RVT = 0.44; Fig. 3a, c).

**Figure 3:**
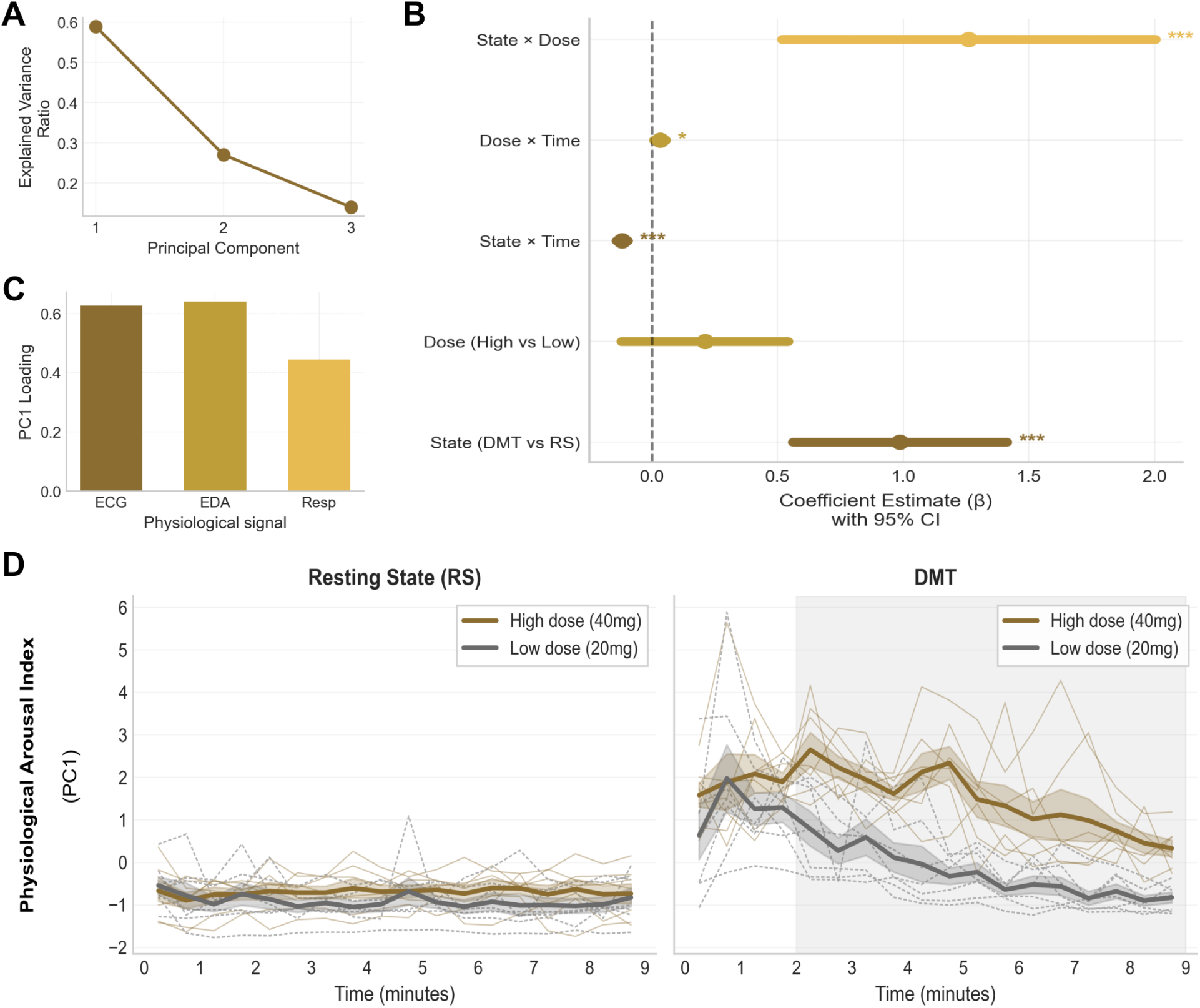
Consistent autonomic activation captured by a single latent component. **a**, Scree plot showing the explained variance ratio for Principal Components (PCs) derived from joint physiological data. PC1 explains 58.9% of the total variance across the three signals. **b**, Fixed-effect coefficients (*β* and 95% CI) from the LME model fitted on the Physiological Arousal Index (PC1 scores). Asterisks indicate significant terms (* *p <* .05, *** *p <* .001; FDR-corrected). **c**, PC1 loadings, all positive: ECG (HR), 0.63; EDA (SMNA), 0.64; Respiration (RVT), 0.44. **d**, Physiological Arousal Index time courses (mean s.e.m.; *n* = 7) during Resting State (RS) and DMT; thin lines show each participant (High dose in colour, Low dose in grey, dashed). Shaded vertical bands indicate clusters of significant High vs. Low differences (cluster-based sign-flip permutation tests; one-tailed for DMT, two-tailed for RS; see Methods). Source data are provided as a Source Data file (Supplementary Data 1).

In the subsample of seven participants with concurrent recordings across all three modalities, the LME model on this index revealed a State × Dose interaction, *β* = 1.26, 95% CI [0.52, 2.00], *p*_FDR_ *<* .001, indicating that systemic autonomic arousal was higher in the High dose condition specifically during the DMT state (Fig. 3b). This effect was temporally dynamic (State × Time: *β* = − 0.12, *p*_FDR_ *<* .001), with the group-mean index peaking 2.0–2.5 min after inhalation under the High dose and the separation between doses largest at approximately 4–5 min post-inhalation. The cluster-based permutation test identified a single sustained cluster of High *>* Low separation spanning approximately 2–9 min post-inhalation (cluster mass = 52.9, exact *p* = .016, *n* = 7; Fig. 3d). This epoch encompasses the dose-separation cluster observed for SMNA and overlaps most of that observed for RVT, consistent with the composite index integrating the temporally distinct contributions of its constituent modalities. When the same analysis was recomputed over the full extended window (0–19 min, 38 windows), dose separation formed three clusters spanning approximately 1.5–10, 10.5–14.5 and 16–19 min post-inhalation (exact *p* = .023 each), indicating that autonomic dose separation persists throughout the recorded session (exploratory, Supplementary Fig. 5). Individual participant trajectories are shown in Supplementary Fig. 6.

### Affective dynamics

To characterise the subjective emotional experience, we analysed time-resolved retrospective ratings obtained with the Temporal Experience Tracing (TET) method. Retrospective ratings of subjective experience (Fig. 4a) revealed two dominant latent dimensions accounting for 72.8% of the total variance (Fig. 4d). The first component (PC1; 41.0% variance) reflected general Arousal, loading positively on Emotional Intensity, Interoception, Anxiety, and Unpleasantness (Fig. 4e). The second component (PC2; 31.8% variance) captured Valence, exhibiting a bipolar structure with positive loadings on Pleasantness and Bliss, and negative loadings on Unpleasantness and Anxiety.

**Figure 4:**
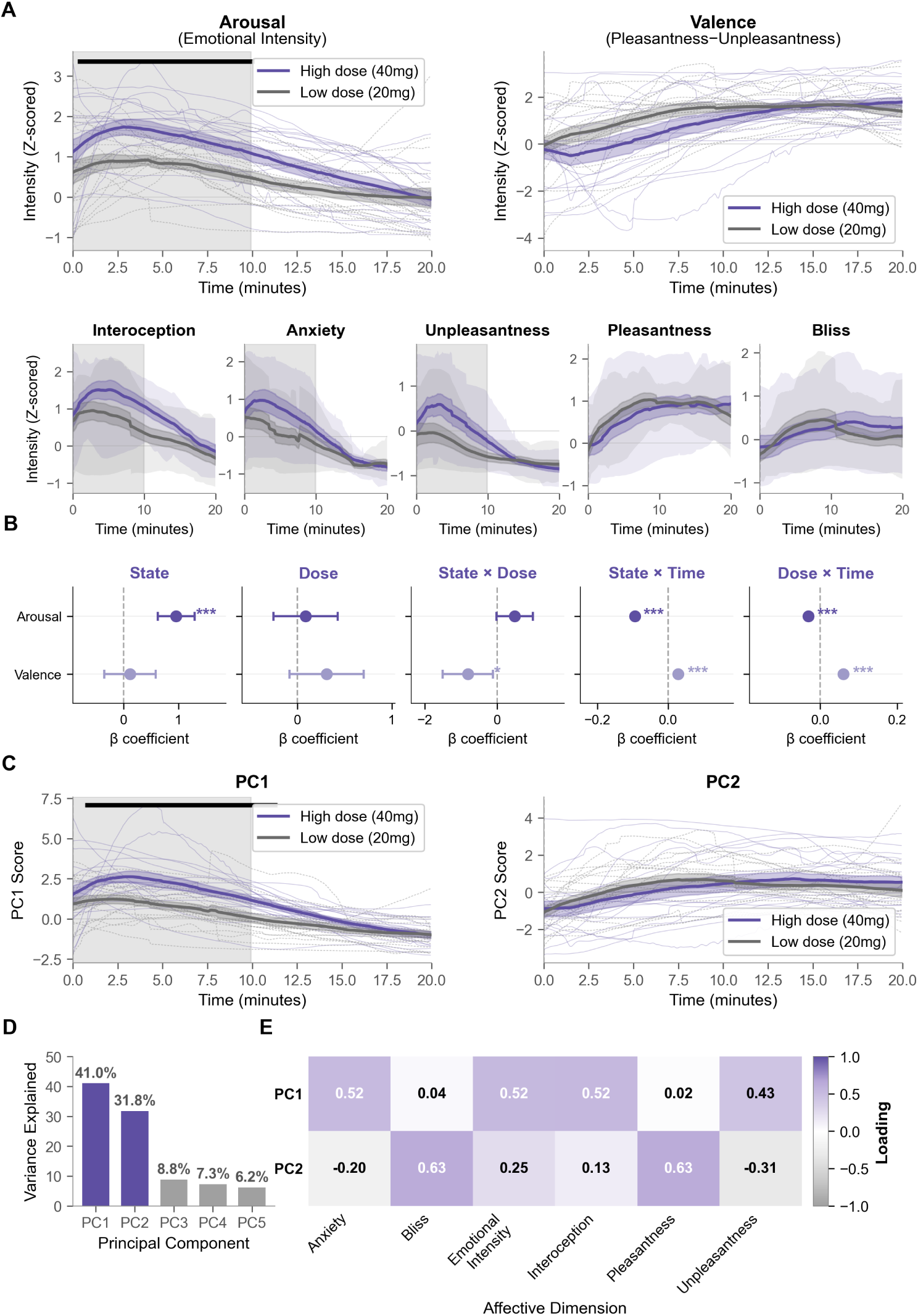
Two latent dimensions characterise the dose-dependent affective dynamics of DMT. **a**, Time courses of retrospective ratings (*z*-scored; mean s.e.m.; *n* = 18). Top: Affective Arousal Index (Emotional Intensity) and Affective Valence Index (Pleasantness minus Unpleasantness); thin lines show each participant (High dose in colour, Low dose in grey, dashed). Bottom: Individual affective items; the narrow band gives the mean s.e.m. and the wide band the 10th–90th percentile across participants for each dose. Grey background shading indicates clusters with a significant DMT vs. RS difference; black horizontal bars indicate significant dose-effect clusters within DMT (High vs. Low; cluster-based sign-flip permutation tests, two-tailed; see Methods). **b**, Fixed-effect coefficients (*β* and 95% CI) from LME models. Asterisks indicate significant terms (* *p <* .05, *** *p <* .001; FDR-corrected). **c**, Time courses of PC1 and PC2 scores (mean s.e.m.; thin lines show each participant); shading and bars are as in **a**. **d**, Scree plot showing that the first two principal components explain 72.8% of the total variance. **e**, PCA loadings heatmap for PC1 and PC2. Source data are provided as a Source Data file (Supplementary Data 1).

LME models on the Affective Valence Index revealed a significant State *×* Dose interaction, *β* = *−*0.81, 95% CI [*−*1.50*, −*0.12], *p*_FDR_ = .022, reflecting a dose-dependent decrease in valence (shift towards difficult/challenging content) during DMT relative to rest (Fig. 4b). For the Affective Arousal Index (Emotional Intensity), the State *×* Dose interaction, averaged over the recording, was positive but its confidence interval narrowly included zero, *β* = 0.49, 95% CI [ −0.01, 0.99], *p*_FDR_ = .056. Cluster-based permutation testing nevertheless localised a dose separation to a single sustained epoch spanning approximately 0.3–10 min post-inhalation (exact *p* = .025; Fig. 4a, black bar), paralleling the sustained autonomic dose separation described above. Within the DMT state the two components followed opposing trajectories (Fig. 4c): under the High dose, PC1 peaked at approximately 3 min post-inhalation and declined thereafter, whereas PC2 was at its most negative immediately after inhalation, crossed zero at approximately 6 min and reached its maximum at approximately 14 min. The cluster-based permutation test localised the dose difference in PC1 to approximately 0.7–11.4 min post-inhalation (cluster mass = 448.5, exact *p* = .021), with no dose cluster for PC2. Quantifying the opposing temporal profiles of arousal and valence, the PC1 and PC2 time courses, averaged across participants and doses, were strongly negatively correlated across the DMT state (Pearson *r* = − .64). Among the individual dimensions, Unpleasantness (*β* = 0.62, 95% CI [0.23, 1.01], *p*_FDR_ = .002) and Interoception (*β* = 0.50, 95% CI [0.02, 0.98], *p*_FDR_ = .041) showed significant State × Dose interactions; the estimate for Anxiety (*β* = 0.42, *p*_FDR_ = .081) was in the same direction but not significant, and those for Pleasantness and Bliss were close to zero (Fig. 4a, bottom panels). The State × Dose estimates for the affective measures were unchanged when TET ratings were downsampled to 20-s and 30-s bins (Supplementary Table 2). The dose-separation cluster for Emotional Intensity was virtually identical at 4-s, 20-s, and 30-s resolution (0.3–10.1, 0.3–10.0 and 0.5–9.5 min; exact *p* = .025–.027).

### Coupling between physiology and subjective experience

Having characterised physiological and affective responses separately, we next asked whether autonomic activation tracked the subjective experience. All coupling analyses in this section were computed on the intersection subsample with concurrent valid recordings across HR, SMNA, and RVT (*n* = 7). We first examined linear associations between individual physiological markers and affective dimensions. During the DMT state, Emotional Intensity correlated positively with all autonomic measures (HR: *r* = .44; SMNA: *r* = .36; RVT: *r* = .43; all *p*_FDR_ *<* .001). Conversely, the Affective Valence Index showed a weak negative correlation with HR (*r* = − .15, *p*_FDR_ = .045). Examination of the Resting State also revealed significant associations between individual physiological measures and affective dimensions, most prominently between HR and the Affective Valence Index (*r* = − .35, *p*_FDR_ *<* .001), Anxiety (*r* = .24, *p*_FDR_ *<* .001), and Unpleasantness (*r* = .22, *p*_FDR_ = .002), as well as between RVT and Anxiety (*r* = .32, *p*_FDR_ *<* .001). Importantly, Emotional Intensity did not show significant associations with any physiological measure during rest (all *p*_FDR_ *>* .05), indicating that the coupling between autonomic arousal and subjective intensity was specific to the DMT state.

To quantify the coupling between the composite Physiological Arousal Index (PC1) and subjective experience, we regressed each TET dimension onto PC1, separately by*_√_*<u>stat</u>e (Supplementary Fig. 7). The Pearson correlation between PC1 and each TET dimension equals *r* = sign(*β*^^^) *· R*^2^. Crucially, the relationship was state-dependent. During the Resting State, the Physiological Arousal Index did not correlate with Emotional Intensity (*r* = .07, *R*^2^ = .004, *β* = −0.04, *p* = .294). In contrast, during DMT, the Physiological Arousal Index showed a moderate-to-strong correlation with Emotional Intensity (*r* = .55, *R*^2^ = .30, *β* = 0.32, 95% CI [0.26, 0.38], *p <* .001), indicating that autonomic activation and subjective arousal covary during the DMT state. The analysis is correlational, so it does not establish the direction of this association. We return to this point in the Discussion. For valence-related dimensions, higher physiological arousal was associated with lower Valence (more unpleasantness) in both states, though these correlations were small (Rest: *r* = .25, *R*^2^ = .06, *β* = −0.34, *p <* .001; DMT: *r* = .15, *R*^2^ = .02, *β* = −0.15, *p* = .021). Beyond Emotional Intensity and the Affective Valence Index, the Physiological Arousal Index also showed weaker but significant associations with individual affective dimensions: during DMT, higher Physiological Arousal Index scores were associated with greater Unpleasantness (*β* = 0.13, *R*^2^ = .06, *p <* .001) and Interoception (*β* = 0.13, *R*^2^ = .04, *p* = .002), whereas during the Resting State higher scores were associated with lower Pleasantness (*β* = −0.23, *R*^2^ = .04, *p* = .001) and lower Interoception (*β* = −0.18, *R*^2^ = .04, *p* = .002); Bliss and Anxiety showed no significant association with the Physiological Arousal Index in either state.

### Multivariate associations between autonomic signals and affective experience

Finally, to identify shared latent dimensions connecting the full physiological and affective spaces, we performed Canonical Correlation Analysis (CCA), which revealed markedly different structures across states (Fig. 5).

**Figure 5:**
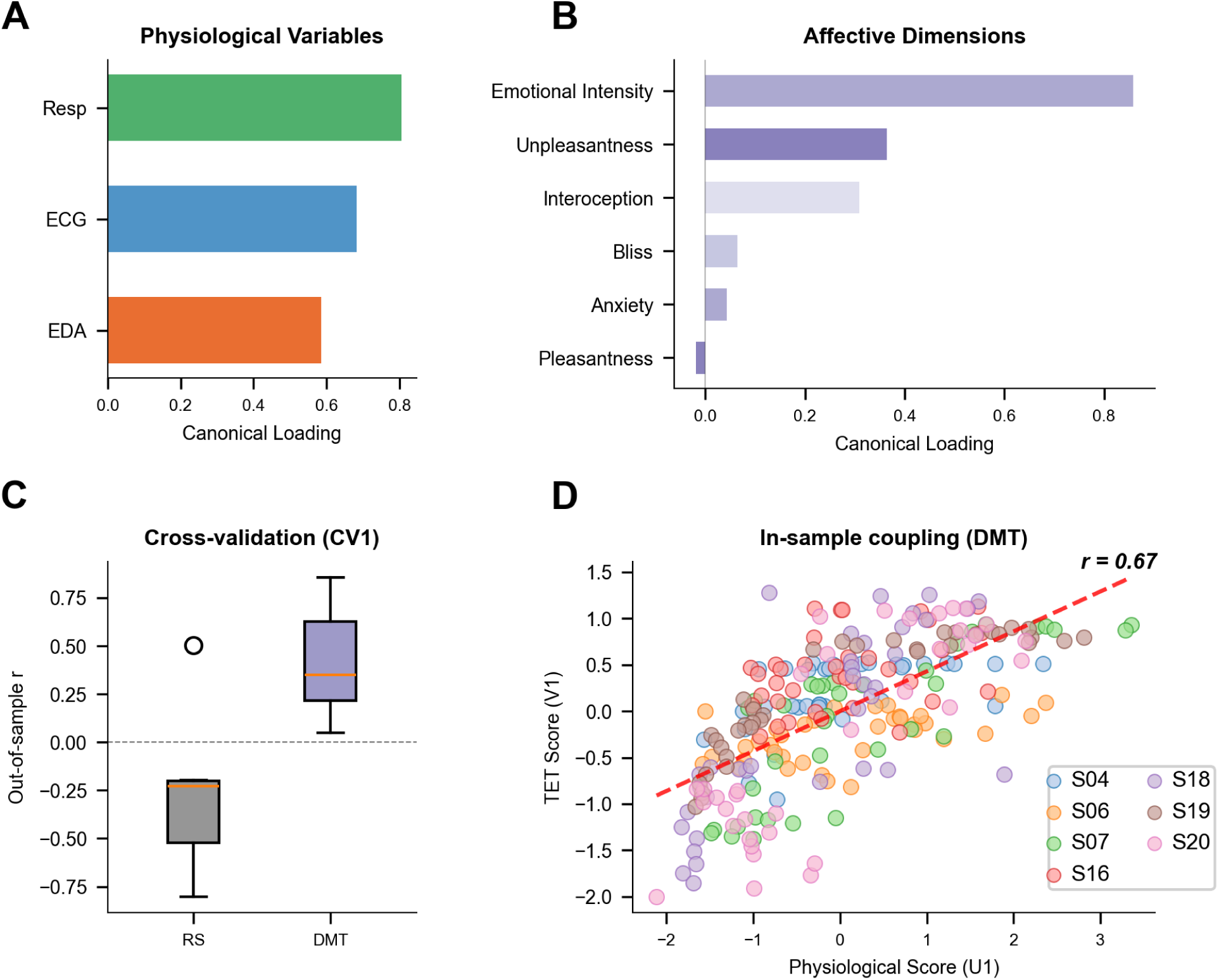
Canonical correlation analysis reveals physiological–affective coupling specific to DMT. **a**, **b**, Canonical loadings on the first canonical variate (CV1) during DMT. Physiological variables (**a**) all loaded positively: Resp (RVT), *r* = .81; ECG (HR), *r* = .68; EDA (SMNA), *r* = .59. Of the affective dimensions (**b**), Emotional Intensity (*r* = .86), Unpleasantness (*r* = .36) and Interoception (*r* = .31) exceeded the substantive loading threshold (*|r| >* .30; see Methods). **c**, Leave-one-subject-out cross-validation performance (*n* = 7). Box plots display out-of-sample correlations (*r*_oos_) across the seven folds during DMT (mean *r*_oos_ = .42, *p* = .010) and Resting State (RS; mean *r*_oos_ = .28, n.s.); the centre line is the median, the box spans the interquartile range and the whiskers extend to the most extreme fold within 1.5 times the interquartile range from the box; folds beyond this are shown individually. **d**, In-sample scatterplot of physiological (U1) versus affective (V1) canonical scores (30-s bins) during DMT. Points are coloured by participant; the regression line shows the linear fit (*r* = .67). Source data are provided as a Source Data file (Supplementary Data 1).

During the DMT state (*n* = 7), a “General Arousal” mode linked systemic autonomic activation (RVT: *r* = .81; HR: *r* = .68; SMNA: *r* = .59) to Emotional Intensity (*r* = .86), with smaller loadings for Unpleasantness (*r* = .36) and Interoception (*r* = .31; Fig. 5a–b), yielding an in-sample canonical correlation of *r* = .67. Although the permutation test did not reach significance (*p*_PERM_ = .118), leave-one-subject-out cross-validation yielded above-chance out-of-sample correlations (mean *r*_oos_ = .42, *p* = .010; Fig. 5c), supporting the generalisability of this physiological–affective pattern across participants. These findings should be interpreted as hypothesis-generating pending replication in larger samples.

In the Resting State, the first canonical mode (in-sample canonical correlation *r* = .56, *p*_PERM_ = .494) linked heart rate (*r* = .94) to Anxiety (*r* = .61) and Unpleasantness (*r* = .48); however, both the non-significant permutation test and the negative out-of-sample correlations (mean *r*_oos_ = − .28; Fig. 5c) indicate that this structure is weak and idiosyncratic, failing to generalise across participants.

## Discussion

The acute autonomic effects of DMT took the form of a transient state of heightened arousal in which, in most participants receiving the High dose, heart rate exceeded what vagal withdrawal alone can produce, indicating active sympathetic engagement. This interpretation is supported by state- and dose-dependent effects of DMT on cardiac, electrodermal, and respiratory measures. Integrating these signals into a single multimodal index of autonomic arousal revealed significant time-resolved differences between doses sustained from approximately 2 min after inhalation to the end of the analysis window, which tracked the subjective affective dynamics assessed using the TET method.

Our findings are consistent with previous studies demonstrating that serotonergic psychedelics modulate markers of ANS activation, including increases in HR, blood pressure, and pupil diameter^28–30^. In the case of intravenous DMT, prior work has shown a pronounced initial increase of HR, followed by a gradual return to baseline typically completed 20 min after injection^31^. After a bolus, heart rate, blood pressure, and subjective intensity all peak within about 2 min, in line with the rapid early pharmacokinetics of DMT^26,28^. Notably, when DMT is administered using a bolus injection followed by constant-rate infusion, subjective effects can be prolonged without corresponding increases in HR beyond the initial minutes of the experience^46^. Together with the present findings, this dissociation suggests that DMT-induced autonomic activation is predominantly linked to the onset of the experience—potentially reflecting an acute emotional or salience-related response—rather than tracking serotonergic receptor occupancy over time.

Our results further indicate that both doses of DMT produced a rapid increase in HR following inhalation; however, these effects were more sustained than those reported in previous studies using intravenous administration, with HR remaining significantly elevated for the High dose relative to the Low dose even approximately 9 min post-inhalation^46^. Because the present study was conducted in naturalistic settings, this prolonged elevation cannot be attributed solely to differences in administration route. Instead, it may reflect interactions between pharmacokinetic factors, contextual variables (i.e., set and setting), and the subjective intensity of the experience^47^.

An intriguing aspect of our findings is that dose-dependent physiological divergence followed modality-specific temporal profiles. Heart rate and respiratory measures distinguished between doses from the earliest phase of the experience (emerging within the first 2 min post-inhalation), consistent with the rapid onset of subjective effects and with the observation that emotional arousal ratings already differed between doses during this early window. In contrast, electrodermal activity—a more specific marker of sympathetic efferent output—showed dose-dependent differences beginning only at approximately 4 min, forming a single cluster at 4–5.5 min post-inhalation. This asymmetry may reflect distinct temporal dynamics across autonomic effectors. Heart rate and respiratory measures can be modulated on short timescales, rapidly tracking the onset of the DMT experience in line with the fast temporal profile of its subjective and neural effects^7,9,45^. Sudomotor responses, mediated exclusively by sympathetic cholinergic fibres, may instead exhibit slower dynamics or reach a transient ceiling immediately following drug onset, attenuating dose-dependent differences at peak intensity^48^. Under this interpretation, dose effects on sudomotor output emerge primarily after the peak of the experience, reflecting a delayed dose-dependent sympathetic engagement—consistent with the observed divergence in electrodermal activity beginning at approximately 4 min and with sustained differences in subjective emotional arousal during the first ten minutes of the experience.

While sympathetic modulation induced by psychedelics is well established, their effects on parasympathetic activity remain less clear. Olbrich and colleagues examined HRV during the acute effects of LSD and ketanserin, a 5-HT_2A_ receptor antagonist, reporting evidence for increased sympathetic activity under LSD, with ketanserin attenuating these effects and concomitantly increasing indices interpreted as parasympathetic tone^29^. In the same study, sympathetic activity was positively associated with the intensity of subjective effects, whereas parasympathetic indices showed the opposite relationship. Bonnelle and colleagues investigated the interplay between sympathetic and parasympathetic influences during the acute effects of DMT, reporting an initial sympathetic-dominant stress response during approximately the first 8–11 min post-injection, followed by apparent sympathovagal coactivation emerging during the subsequent core experience window as vagal tone rebounded^31^. However, these interpretations are limited by the reliance on cardiac measures alone and the absence of complementary physiological indices with greater specificity for sympathetic efferent activity^32^. In contrast, our simultaneous assessment of ECG and EDA provides convergent evidence that DMT acutely increases SMNA, a selective marker of sympathetic activation^49,50^. These findings indicate that the observed increases in heart rate cannot be attributed solely to parasympathetic (i.e., vagal) withdrawal and involve active sympathetic efferent engagement during the onset of the DMT experience. This inference does not rest on the electrodermal evidence alone. Because vagal withdrawal alone can raise heart rate at most to the intrinsic rate of the sinoatrial node, and under the High dose peak heart rate exceeded the estimated intrinsic rate in most participants and on average across the sample, the rise indicates sympathetic drive in addition to any vagal withdrawal. Time-domain variability indices were also examined (Supplementary Fig. 4). They declined under DMT in a dose-dependent manner, but the decline could not be separated from the accompanying shortening of the cardiac cycle, which is why the argument above rests on the heart rate attained rather than on variability.

We observed covariance across the recorded physiological signals, strongest between HR and SMNA. Data-driven dimensionality reduction yielded a Physiological Arousal Index that accounted for approximately 59% of the total variance, with positive contributions from all three modalities. This index exhibited significant state-by-dose interactions, reflecting dose-dependent differences that emerged after DMT inhalation, as well as sustained divergence between doses during the post-peak phase of the experience. Moreover, the Physiological Arousal Index showed robust associations with the temporal dynamics of subjective reports. Together, these findings suggest that peripheral monitoring of DMT-induced affective states—such as in experimental or clinical contexts—may benefit from incorporating electrodermal and respiratory measures alongside cardiac recordings, rather than relying on ECG alone, consistent with the recommendation that autonomic assessment combine several complementary tests rather than rely on any single one^51^. In particular, the inclusion of electrodermal activity provides a time- and cost-effective enhancement, given its specificity for sympathetic activation.

Previous studies have linked the intensity of psychedelic experiences to markers of ANS activation; however, subjective effects in these studies were typically assessed retrospectively using standard psychometric questionnaires (e.g., the 5D-ASC or 11D-ASC), which limits their utility for relating physiological measures to the temporal evolution of affective states^35^. We addressed this limitation by incorporating subjective reports obtained using the TET method, a phenomenological approach originally developed for the study of meditative states^39,40^ and later applied to high-ventilation breathwork^44^ and DMT-induced altered states of consciousness^10^. Although TET captures multiple experiential dimensions relevant to psychedelic states, it does not explicitly include dimensions corresponding to affective valence and arousal. We therefore adopted a data-driven approach to characterise affective dynamics, identifying two dominant latent components within the multidimensional TET reports. The first component was associated with emotional arousal, exhibiting positive loadings on Emotional Intensity, Interoception, Anxiety, and Unpleasantness. The second component reflected emotional valence, with positive loadings on Pleasantness and Bliss and negative loadings on Unpleasantness and Anxiety.

These two components displayed biphasic and temporally opposed dynamics, with the PC1 and PC2 time courses, averaged across participants and doses, showing a strong negative correlation during the DMT state (*r* = − .64). Under the High dose the arousal component peaked approximately 3 min after inhalation and returned gradually towards baseline over the remainder of the recording. The Physiological Arousal Index peaked earlier, within the first 2.5 min, although its separation between doses was largest between 4 and 5 min. Given the difference in temporal resolution between a retrospective trace and a 30-s physiological window, we do not read this ordering as evidence that either measure leads the other, a point we return to below. In contrast, the valence component initially shifted towards negative values before transitioning towards positive values later in the experience. Consistent with previous findings^26^, this pattern suggests an early phase characterised by heightened anxiety, followed by a more sustained period dominated by feelings of pleasantness and bliss. Notably, the High dose increased the negative affective tone of the experience: the Affective Valence Index shifted towards more negative values during DMT (State × Dose: *β* = − 0.81, *p*_FDR_ = .022), and Unpleasantness and Interoception were the individual dimensions showing a significant dose effect (*β* = 0.62 and *β* = 0.50), indicating that higher doses intensified the challenging and bodily content of the experience. Anxiety, in contrast, was elevated under DMT relative to rest (Fig. 4a) but did not differ significantly between doses, in either the LME model or the cluster-based test. This is consistent with anxiety at the onset of the DMT state reflecting a response to the sudden alteration of conscious state produced by the drug rather than simply the overall intensity of subjective effects.

We observed consistent associations between subjective reports of affective experience and both uniand multimodal measures of autonomic arousal. Specifically, during the acute effects of DMT, Emotional Intensity (the Affective Arousal Index) showed significant correlations of comparable magnitude with HR, SMNA, and RVT. In contrast, the Affective Valence Index exhibited a weaker and oppositely directed association, limited to HR. This contrast suggests that autonomic activation covaries more closely with the intensity of affective experience than with its valence, whose relationship to physiology may be more indirect or mediated by additional factors. This pattern supports the link between emotional arousal and sympathetic activation, reflected in the simultaneous increase of these physiological measures^20,21^. An exploratory canonical correlation analysis yielded a convergent pattern specifically during the DMT state, with all autonomic measures showing associations with Emotional Intensity—a finding that, as expected given the reduced affective and physiological variability at rest, did not generalise to the Resting State. The relationship between emotional intensity and physiological indices of sympathetic activation has been consistently reported across previous studies; however, the capacity of peripheral physiological signals to predict affective valence—or to distinguish between specific emotional states—remains less clear^23,52^. The present analyses are correlational, and we did not attempt to model the direction of the physiology–experience coupling. With 18 matched 30-s windows per participant, state and dose, the time series are likely too short to identify a directional model for a single participant^53^, and pooling across participants would require a model of how those dynamics vary from one participant to the next^54^. A finer temporal resolution would provide more points but meets two limits of the measurements. The TET trace is drawn retrospectively^10,40^, anchored by chimes every two minutes, and its precision on the time axis has not been quantified. This uncertainty barely affects correlations computed over several minutes, but it is of the same order as the lags a directional model would estimate. Sudomotor nerve activity is recovered from skin conductance by deconvolution with an assumed impulse response^55^, although the shape of the skin conductance response varies between individuals^56^, so at fine resolution its timing partly reflects the assumed response rather than the underlying neural drive. The same limit applies to directional analyses between the physiological signals themselves. Directional inference would therefore require a phenomenological measure of known temporal precision and a participant-specific model of the sweat-gland response.

When generalised to other psychedelic compounds and clinical populations, our results may contribute to the development of physiological biomarkers of psychedelic-induced affective states^57^. Given the relationship between clinical outcomes and the acute subjective effects induced by psychedelics, the identification of reliable objective markers of affective states could facilitate improved quantification of treatment efficacy^58^. Previous work has suggested that features of sympathetic and parasympathetic activation can predict spiritual and insightful aspects of the experience, which are relevant for outcome prediction^31^. This line of research could be expanded by adopting the multimodal characterisation introduced in our study. Furthermore, the observed link between sympathetic arousal and negative affective tone raises the question of whether targeted interventions, such as slow-paced breathwork during the come-up phase, could dampen physiological arousal^59^ and thereby shift the affective balance towards more positively valenced experiences, with potential relevance for clinical contexts. This hypothesis warrants direct empirical investigation. However, given evidence that the intensity of the psychedelic experience is positively associated with clinical outcomes^60^, interventions aimed at reducing arousal should be carefully evaluated to ensure that any attenuation of challenging affect does not also reduce therapeutic efficacy. Future work should also examine the effects of longer-lasting serotonergic psychedelics (such as psilocybin or LSD) on multimodal measures of autonomic arousal, and assess whether fluctuations in these measures track changes in affective states under conditions of relatively stable plasma drug concentrations, in contrast to the rapid pharmacokinetics characteristic of DMT.

The present study has several limitations, some of which are inherent to investigations of psychedelic experiences conducted in naturalistic settings. Although our design allowed participants and experimenters to remain blinded to dose, the facilitator who assisted participants during DMT administration was not blinded to dose assignment. Consequently, unintentional behavioural cues from the facilitator cannot be fully excluded as a potential influence on participants’ experience. In addition, although the rate at which participants identified the dose received did not differ between dose conditions, identification was above chance overall, indicating that perceptual blinding was incomplete; consequently, contributions of subjective expectancy to the observed dose-related effects cannot be entirely excluded. In addition, estimates of DMT dose were derived from the mass of freebase material consumed, rather than from direct pharmacokinetic measurements. As a result, dose estimates may be imprecise due to variability in extraction efficiency. While inhalation represents a route of administration with high ecological validity for DMT, it also introduces variability in effective dosing related to individual differences in inhalation technique and pulmonary absorption. Dose delivery by inhalation is therefore less precise than intravenous (or intramuscular) injection, in which the administered amount is known rather than estimated. Because the studies with which we compare our time courses administered DMT intravenously^7,26,28,31,46^, differences in the magnitude and timing of effects between those studies and ours may partly reflect the route of administration. This additional source of variability is, however, expected to attenuate rather than inflate dose-related contrasts: imprecision in the delivered dose adds noise to the High versus Low comparison, so the dose-dependent effects reported here are likely to be conservative estimates of those obtainable under pharmacokinetically controlled administration. Although the TET method provides time-resolved phenomenological data, it relies on retrospective reconstruction of experience; during the acute peak of DMT (particularly at the higher dose) participants may experience pronounced time distortion and disrupted episodic encoding, which could limit the temporal accuracy of retrospective traces during this phase. The sample was predominantly male, which limits the generalisability of our findings given established sex differences in autonomic reactivity^50,61^ and psychedelic-induced emotional experience^62^. The comparison with the intrinsic rate of the sinoatrial node also carries caveats of its own. The regression we applied was derived under combined pharmacological blockade in a separate population^63^ and predicts an individual’s rate only approximately, since individuals of the same age differ under blockade^36,37^, and intrinsic rate is itself lowered by aerobic conditioning^64^, which we did not assess. We therefore report the counts against the upper 95% limit of the estimate as well as against the estimate itself, and the comparison should be read as a limit on the vagal contribution across the sample rather than as an exact classification of each participant. Furthermore, because pharmacokinetic measurements were not obtained, we cannot exclude the possibility that the observed associations between autonomic and affective measures partly reflect shared pharmacokinetic variation—both covarying with plasma DMT concentration over time—rather than direct physiological–affective coupling. Future studies incorporating plasma DMT measurements will be needed to disentangle these contributions. Finally, the sample size was limited and was further reduced for multimodal analyses requiring high-quality recordings across all physiological signals, which should be treated as preliminary and hypothesis-generating. Future studies with larger and more diverse samples, conducted under both naturalistic and controlled laboratory conditions, will be important for confirming the present findings and for further disentangling the contributions of set and setting to DMT-induced affective states and their physiological correlates.

In conclusion, inhaled DMT produces a dose-dependent autonomic activation that, in most participants receiving the High dose, raises heart rate above the rate attainable by vagal withdrawal alone, indicating active sympathetic engagement rather than the release of parasympathetic restraint alone. The three effectors separate the doses at different times: cardiac and respiratory measures within the first two minutes, whereas electrodermal activity, the most specific of the three for sympathetic efferent output, only later in the experience. This activation covaries in time with the intensity of the affective experience, while affective valence follows a distinct trajectory. Beyond DMT, the approach shows that an active sympathetic contribution to a psychedelic-induced cardiovascular response can be inferred using measures that are inexpensive and portable enough for naturalistic settings, which is a prerequisite for developing physiological markers of psychedelic-induced emotional processes.

## Methods

### Participants

Nineteen volunteers (17 self-identified as male, 2 as female; age *M* = 33.9, *SD* = 4.8 years) were enrolled following community recruitment and social media announcements. Sex and gender were collected by self-report. Given the strong sex imbalance, which reflects the demographics of the recruited community, sex was not included as a factor in the primary analyses. Eligibility was determined via a non-diagnostic interview with a mental health professional, based on the Structured Clinical Interview for DSM-IV, Clinical Trials Version^65^ and standard safety guidelines for human psychedelics research^66^. Inclusion criteria comprised being aged 21–65, having had at least two prior experiences with psychedelics, and having abstained from psychoactive substances, including alcohol, caffeine, and tobacco, for at least 24 hours before each session. Prior exposure to DMT itself varied widely across participants and was positively skewed (*M* = 4.6, *SD* = 5.5, *Mdn* = 2, range 0–20 occasions); three participants had no prior DMT experience and met the eligibility criterion through prior experience with other psychedelics. Exclusion criteria included personal DSM-IV diagnoses of psychotic or bipolar disorders, a first- or second-degree family history of these conditions, substance abuse or dependence in the previous five years (excluding nicotine), current or recurrent depressive or anxiety disorders, eating disorders, neurological illness, pregnancy, trait anxiety more than one standard deviation above normative means as measured by the State–Trait Anxiety Inventory^67^, and ongoing psychiatric medication. Participants who had previously experienced adverse effects of psychedelic substances, such as lasting psychological difficulties or risky behaviours, were also excluded. All participants provided written informed consent. The study complied with the Declaration of Helsinki and was approved by the Research Ethics Committee at José María Ramos Mejía General Hospital in Buenos Aires, Argentina, with code CE000005.

### Study design

We used a within-participant, randomised and counterbalanced 2 × 2 design under naturalistic conditions, with factors Dose (20 mg vs. 40 mg freebase DMT; between sessions) and State (pre-dose resting state [RS] vs. DMT acute effects within session). Each participant completed two experimental sessions on separate days, spaced by at least two weeks. On each session day, participants first completed the eyes-closed resting state (RS) baseline and then performed a single prolonged inhalation of freebase DMT at the assigned dose. Sessions took place in a quiet, comfortable room. During recordings, participants rested in a reclined position with eyes closed under an eye mask. Dose order was counterbalanced across participants. Dose identity (20 vs. 40 mg) was masked from the participants until data acquisition for both sessions had concluded. The facilitator who assisted the participants during the sessions prepared and administered both doses and was therefore not blinded to dose identity. All other personnel present during the recording sessions were blinded to the dose administered in each session. Blinding was not maintained during data analysis, as dose was an experimental factor in the statistical models. After both sessions had been completed, participants were asked to indicate which dose (20 vs. 40 mg) they believed they had received in each session. Blinding efficacy was assessed with two chi-square tests on the resulting 38 sessions: a one-sample goodness-of-fit test against chance (50%) and a 2×2 test of independence between administered dose and identification accuracy. Participants correctly identified the dose in 26 of 38 sessions (12/19 [63%] for 40 mg, 14/19 [74%] for 20 mg); identification was above chance overall (*χ*^2^(1*, N* = 38) = 5.16, *p* = .023) but did not differ between dose conditions (*χ*^2^(1*, N* = 38) = 0.49, *p* = .485), indicating that subjective discriminability of the two doses was comparable.

### Procedure

On each dosing day, participants first completed a 10-min eyes-closed RS baseline with continuous physiological recording. Immediately afterwards, they completed retrospective temporal traces of subjective experience for the RS baseline. Participants then performed a single prolonged inhalation of freebase DMT at the assigned dose. For all time-locked analyses, *t*_0_ was defined separately for each state: for RS, *t*_0_ was the onset of the baseline recording; for DMT, *t*_0_ was the end of the DMT exhalation. After inhalation, participants remained reclined with eyes closed under an eye mask for a further 20 min while recordings continued. Throughout the RS and post-inhalation periods, auditory chimes every two minutes served as temporal anchors to aid later retrospective tracing of the experience. After the 20-min period, and once ready, participants completed their DMT retrospective traces, followed by a brief chronological debrief interview (audio recorded for documentation but not analysed in the present study).

### Affective phenomenology

Subjective dynamics were assessed using Temporal Experience Tracing (TET), a retrospective, time-resolved quantitative phenomenological method^10,44^. After each recording block, and separately for each predefined experiential dimension, participants drew a continuous curve on a two-dimensional canvas in a mobile application (Human Experience Dynamics Ltd), where the horizontal axis represented elapsed session time (10 min for RS, 20 min for DMT) and the vertical axis represented the perceived intensity of that dimension over the course of the block. The trace for each dimension was completed at the participant’s own pace and was not time-locked to real-time playback of the experience. Before the first session, participants received standardised instruction and practice on the application, and practice traces were reviewed to ensure procedural understanding. The fifteen dimensions were translated into Spanish and selected *a priori* based on literature specifically defining experiential constructs relevant to DMT^10^. For the statistical analyses reported here, we focused on a predefined affective/autonomic subset that captures emotional and bodily aspects of the experience: Pleasantness, Unpleasantness, Emotional Intensity, Interoception, Bliss, and Anxiety. Expanded definitions of all dimensions are provided in Lewis-Healey *et al.*^10^.

### Subjective data processing and dimensionality reduction

Auditory chimes delivered every two minutes, presented at two distinct pitches to demarcate the first and second halves of the session, served as temporal anchors in the TET interface to guide the retrospective placement of experiential features. TET time series were sampled at a native resolution of 0.25 Hz, yielding 150 samples for the 10-min RS baseline and 300 samples for the 20-min DMT period. To control for inter-individual differences in scale usage, raw TET values were transformed into within-participant *z*-scores using each participant’s mean and standard deviation calculated across all dimensions and sessions.

To characterise the shared latent structure of the affective TET ratings, principal component analysis (PCA) was applied to the six predefined affective dimensions. Prior to PCA, a second standardisation was performed at the group level (*z*-scoring each dimension across all participants) to ensure comparable loadings. PCA was then conducted on this dataset pooled across participants and sessions. The first two principal components (PC1 and PC2) were retained based on the scree plot, together accounting for 72.8% of the total variance. PC1 loaded positively on Emotional Intensity, Interoception, Anxiety, and Unpleasantness, and was therefore interpreted as reflecting general emotional arousal. PC2 exhibited a bipolar structure, with positive loadings on Pleasantness and Bliss and negative loadings on Unpleasantness and Anxiety, consistent with an affective valence dimension.

Rather than using the PC scores directly as dependent variables, we defined two theory-guided summary indices grounded in the Circumplex Model of Emotion^18^, whose interpretation was corroborated by the PCA structure. For arousal, the original Emotional Intensity dimension was retained, as it was among the dimensions loading most strongly on PC1 and reflects the momentary strength of experience independent of hedonic tone. For valence, a composite Affective Valence Index was computed as the difference between Pleasantness and Unpleasantness (Valence = Pleasantness*_z_* Unpleasantness_z_) at each time point, consistent with the bipolar structure captured by PC2. This approach preserves the interpretability of the original TET dimensions while ensuring that the chosen indices align with the dominant latent structure identified by PCA. For statistical modelling, the native 0.25 Hz time series (i.e., one sample every 4 s) were retained at their original temporal resolution.

### Physiological signals

Peripheral physiology was acquired continuously during both sessions using a Brain Products amplifier with EXG inputs (BrainAmp ExG MR; Supplementary Fig. 1), concurrently with EEG recordings whose acquisition and analysis are reported in Lewis-Healey *et al.*^10^. Electrodermal activity (EDA), electrocardiography (ECG), and respiration were sampled at 250 Hz. ECG was recorded using a modified lead II bipolar montage with electrodes placed below the clavicles, and respiration was monitored via a thoracic effort belt positioned at the level of maximal expansion. All channels were time-locked to state-specific onsets (*t*_0_). Data processing was performed in Python using NeuroKit2^68^, BioSPPy^69^, and custom scripts (https://github.com/tomdamelio/dmt-emotions).

Exclusion criteria were defined *a priori*: a participant was excluded from a given modality if either of the two sessions for that modality contained data losses or non-recoverable artefacts (e.g., excessive movement noise, sensor disconnection, missing data files for early participants S01–S03 where recordings were unavailable). No participant or session was excluded *post hoc* on the basis of the statistical results. Phenomenological data were available for 18 of the 19 participants (one participant excluded due to missing retrospective traces). Final modality-specific sample sizes were *n* = 11 for ECG, *n* = 11 for EDA, and *n* = 12 for respiration. Participant-level exclusion criteria for each physiological measure are detailed in Supplementary Table 3. For multivariate integration analyses requiring simultaneous signals across all three modalities, the sample was restricted to the intersection of these subsets (*n* = 7).

### Electrodermal activity

Raw EDA was decomposed into tonic and phasic components using convex optimisation (cvxEDA;^55^) as implemented in BioSPPy^69^. Following the original cvxEDA formulation, no bandpass filtering was applied prior to decomposition, as the algorithm jointly estimates all signal components and does not require pre-processing of the observed skin conductance signal. The cvxEDA algorithm models the observed skin conductance as the sum of three components: (1) a slow tonic component representing the skin conductance level (SCL), (2) a sparse phasic component representing skin conductance responses (SCRs), and (3) measurement noise. The phasic component is further deconvolved with a biophysically inspired impulse response function to estimate the underlying sudomotor nerve activity (SMNA), which represents the neural drive to the sweat glands. We quantified sympathetic arousal by computing the area under the curve (AUC) of the SMNA signal within non-overlapping 30-s windows aligned to *t*_0_. No baseline correction was applied prior to summarisation.

### Electrocardiography

ECG signals were cleaned using NeuroKit2’s default pipeline, which applies a 0.5 Hz high-pass Butterworth filter (5th-order) followed by powerline removal at 50 Hz. R-peaks were detected using NeuroKit2’s default algorithm, which implements an adaptive threshold approach based on the Pan–Tompkins method^70^. This algorithm applies a series of filters (derivative, squaring, moving window integration) to enhance QRS complexes, followed by adaptive thresholding to identify R-peaks. Inter-beat intervals (IBIs) were computed as the time difference between consecutive R-peaks, and instantaneous heart rate (HR; beats per minute) was derived as HR = 60,000*/*IBI (with IBI in milliseconds). The resulting HR time series was resampled to the original sampling rate (250 Hz) using cubic interpolation. HR traces were visually inspected for artefacts, and values were averaged within non-overlapping 30-s windows over the first 9 min.

R-peaks were corrected with the Kubios algorithm^71^ at preprocessing, for all ECG analyses; the algorithm identifies ectopic, missed, extra, and long/short intervals from the distribution of successive differences, and across the 70 ECG recordings in the dataset 357 of 84,633 beats (0.42%) were flagged and corrected. Because detection errors bias beat-to-beat variability far more than they bias mean rate, an inter-beat interval was additionally treated as invalid for the variability analyses reported in the Supplementary Information if it fell outside 300–2000 ms or deviated by more than 25% from the median of its eleven nearest neighbours, a relative criterion that captures false and missed detections falling within the physiological range. Invalid intervals were discarded rather than interpolated, and successive differences were computed only within runs of consecutive valid intervals, so that no difference spanned a gap. A recording was excluded from the variability analyses if more than 10% of its intervals were invalid; this applied to one recording (participant S16, Low dose, 22.7% invalid, against a median of 0.17% across the 44 recordings of the eleven participants with valid ECG), and that participant was therefore excluded from the variability analyses, leaving *n* = 10. Mean HR was unaffected by this filter (median change across recordings = 0.00 bpm; 0.18 bpm in the excluded recording), so the HR analyses retain the full *n* = 11.

### Heart rate variability

To provide a candidate index of parasympathetic activity, three time-domain variability measures were computed from the corrected inter-beat interval time series: the root mean square of successive differences (RMSSD), the standard deviation of inter-beat intervals (SDNN), and RMSSD divided by the mean inter-beat interval. The last of these addresses the dependence of RMSSD on cycle length: as heart rate rises, intervals shorten and the differences between them shrink proportionally, so part of any RMSSD decrease is arithmetic rather than autonomic^72^. Because variability indices require longer estimation windows than mean rate^73^, and SDNN in particular captures slower fluctuations than a 30-s bin contains^34^, measures were computed in 120-s windows sliding in 30-s steps, a length at which both RMSSD and SDNN have been shown to reproduce the values obtained from 4-to 5-min reference recordings^74^, rather than in the non-overlapping 30-s bins used for the other modalities, and a window was retained only if valid intervals covered at least 80% of its duration and numbered at least 60 (no window was lost under this rule). The ratio of low-to high-frequency spectral power (LF/HF) was not computed, for three reasons. First, the ratio assumes that respiratory variability falls within the conventional high-frequency band^34,75–78^, whereas respiratory frequency in this sample fell below that band in most participants and was often non-periodic during DMT. Second, the rapid rise in heart rate after inhalation violates the stationarity that spectral estimates require^79^. Third, the reading of the ratio as an index of sympathovagal balance has been questioned even when the band assignment holds^80^.

Because the sinoatrial node has an intrinsic firing rate that autonomic blockade reveals^81,82^, vagal withdrawal alone can raise heart rate at most to that rate^36,38^. Each participant’s intrinsic rate was estimated as 118.1 0.57 age^63^ and compared with their peak heart rate under each dose, computed over 60-s windows sliding in 5-s steps within the 9-min analysis window. Resting heart rate was averaged over the 9-min analysis window of the resting-state recording that preceded each dose, the rise was taken as peak minus resting rate, and resting rate, peak rate and rise were compared between doses with two-tailed paired *t*-tests. Because the regression predicts an individual’s rate only approximately, with 95% limits of ±15% for individuals of the same age^37^, counts are also reported against the upper limit of each estimate, the least favourable case for the inference, and the margin between peak and estimated intrinsic rate was tested across participants with one-tailed one-sample *t* and Wilcoxon signed-rank tests.

### Respiration

Respiration signals from the thoracic effort belt were processed using NeuroKit2 with the Khodadad *et al.*^83^ algorithm for peak detection. Respiratory cycles were identified by detecting peaks (inhalation maxima) and troughs (exhalation minima) in the filtered signal. Respiratory volume per time (RVT) was computed following Birn *et al.*^84^ as the product of respiratory amplitude and respiratory rate:

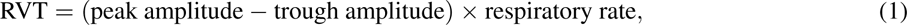

where respiratory rate is the inverse of the breath-to-breath interval. This metric captures both the depth and frequency of breathing, providing an integrated measure of respiratory drive. The RVT time series was visually screened for artefacts and averaged within non-overlapping 30-s windows over 0–9 min.

All physiological metrics were summarised within non-overlapping 30-s windows and then *z*-scored within participant, using the mean and standard deviation of that participant’s window values across all four sessions, to facilitate mixed-effects modelling. Standardising at the level of the analysis window rather than at the level of the raw samples matters here because the three signals differ in how much they fluctuate within a window relative to between windows: the ratio of the sample-level to the window-level standard deviation had a median of 0.98 for HR, 1.57 for RVT and 10.86 for SMNA, the last reflecting the burst-like character of the electrodermal signal. Standardising at the sample level would therefore have made one standard deviation mean a different quantity in each modality, and the coefficients across panels of Fig. 2 would not have been comparable. With window-level standardisation, one unit means the same thing throughout: how far a given window lies from that participant’s average window. Participants missing valid data in any one session for a given modality were excluded from that modality’s analyses.

### Physiological Arousal Index (PC1)

To capture variance shared across autonomic modalities, we performed PCA on the concatenated, within-participant *z*-scored time series of HR, SMNA AUC, and RVT (restricted to the *n* = 7 subsample). The first principal component (PC1) was retained as the Physiological Arousal Index.

### Statistical analyses

All affective and physiological measures were analysed within a linear mixed-effects (LME) framework in Python^85^. For each outcome variable *Y*, the model was specified as:

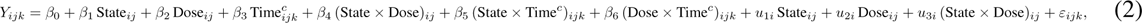

In Python (statsmodels), the fixed-effects part of this model was implemented using the formula: Y State * Dose + Time_c + State:Time_c + Dose:Time_c, with the random-effects structure given below. Here *i* indexes participants, *j* indexes sessions, and *k* indexes the time samples at the native 0.25 Hz resolution (one sample every 4 s). For TET-derived affective measures, the full recording was modelled (RS: 0–10 min, 150 samples; DMT: 0–20 min, 300 samples). For physiological measures, data were summarised into 18 non-overlapping 30 s windows spanning 0–9 min post-*t*_0_. State was coded as RS = 0, DMT = 1; Dose as Low (20 mg) = 0, High (40 mg) = 1. The time index was mean-centred (Time*^c^*) to reduce collinearity between main effects and interactions. Random effects were specified as by-participant slopes for State, Dose, and their interaction ((0 + State + Dose + State:Dose | Subject)), denoted *u*_1_*_i_*, *u*_2_*_i_* and *u*_3_*_i_* in the equation above, so that the structure is maximal with respect to the effects under test^86^. The by-participant slopes were taken as multivariate normal with zero mean and an unstructured covariance matrix. For the physiological outcomes the random intercept was omitted by design: because these outcomes are *z*-scored within participant across that participant’s four sessions, each participant’s mean is exactly zero and their within-participant standard deviation exactly one, so the between-participant intercept variance is zero by construction rather than by estimation (the intraclass correlation is zero for all three physiological measures), and a model with a by-participant intercept alone returns a singular random-effects covariance. The TET dimensions, in contrast, are standardised with a single mean and standard deviation per participant pooled across all dimensions, so each dimension retains a participant-specific mean; the affective models therefore additionally carry a by-participant random intercept. For the physiological outcomes, what varies across participants is the size of the response: the between-participant SD of the State slope ranged from 0.33 (HR) to 0.53 (Physiological Arousal Index), and that of the State × Dose slope from 0.34 (HR) to 1.09 (RVT). Adding the by-participant State × Dose slope improved fit for every physiological outcome, tested against the simplest structure that is estimable here, which carries by-participant slopes for State and Dose only (HR, *χ*^2^(3) = 22.9; SMNA, *χ*^2^(3) = 67.5; RVT, *χ*^2^(3) = 91.7; Physiological Arousal Index, *χ*^2^(3) = 52.3; all *p <* .001); the intercept-only structure is degenerate for the reason just given and yields no likelihood against which to test. Time was entered as a control rather than as a tested effect and does not carry a by-participant slope, since retaining variance components the design does not require costs power^87^. The residual error is *ε_ijk_* (0*, σ*^2^). Models were fit by restricted maximum likelihood (REML), and fixed-effect estimates are reported as standardised coefficients (*β*) with 95% confidence intervals (CI). Effect sizes are reported as Cohen’s *d* for the corresponding within-participant contrast, computed as the mean of that contrast across participants divided by its standard deviation.

To control for multiple testing, Benjamini–Hochberg false discovery rate (BH-FDR) correction^88^ was applied within each model, over three families of fixed effects: the State effect together with its interaction with time, the Dose effect together with its interaction with time, and the State × Dose interaction. The last is a family of one, so for that term the adjusted value equals the raw one. Adjusted *p* values are reported as *p*_FDR_.

The fixed effects of each model were specified in advance, and no alternative sets of fixed effects were compared. To assess the robustness of the main findings to potential violations of the homoscedasticity assumption, each physiological LME was re-estimated with a state-specific residual variance structure (varIdent(form 1 State), R package nlme;^89^).

To characterise the temporal evolution of dose-dependent effects, the LME models were complemented with cluster-based permutation tests comparing High vs. Low dose across time points, computed separately within each state^90^. Where a time course is compared with rest, as for the affective TET measures (Fig. 4) and the heart rate variability measures (Supplementary Fig. 4), the same procedure was applied to DMT vs. RS (two-tailed), after averaging each participant’s two doses, over the time points common to both states. For physiological measures, each time point corresponds to one of the 18 non-overlapping 30 s windows (0–9 min); for affective TET measures, each time point corresponds to one native 4 s sample. Paired differences (High Low) were computed for each participant at each time point and tested against zero with a one-sample *t*-test; time points exceeding the cluster-forming threshold (the *t* value corresponding to an uncorrected *p <* .05 with *n* − 1 degrees of freedom) were grouped into temporally contiguous clusters, and the cluster mass was defined as the sum of *t* values within the cluster. The null distribution of the maximum cluster mass was obtained by exhaustive sign-flip permutation of the participant-level difference time courses (2*^n^* permutations for one-tailed tests and 2*^n−^*^1^ for two-tailed ones, since flipping every sign returns the same statistic), yielding exact cluster-level *p* values corrected for multiple comparisons across time; with *n* = 7 (composite index) the smallest attainable *p* is 1*/*128 ≈ .008. Clusters with *p <* .05 are visualised as shaded regions in the time-course plots. Cluster extent is reported as approximate: the test assigns a single *p* value to each cluster as a whole, and the cluster’s boundaries depend on the uncorrected threshold used to form it and on noise in the windows at its edges, so a significant cluster does not establish that an effect begins or ends exactly where the cluster does^91^. Tests were implemented with MNE-Python^92^. For physiological time courses in the DMT state, one-tailed tests were employed based on the *a priori* directional hypothesis that the High dose would induce greater autonomic arousal (High *>* Low). For TET affective time courses, two-tailed tests were used in both states, given the mixed expected directions across affective dimensions (e.g., Pleasantness could plausibly increase or decrease under the High dose, depending on the character of the individual experience). For the Resting State (RS) physiological measures, where no directional difference was expected, two-tailed tests were also used. For completeness, window-wise paired *t*-tests with Benjamini–Hochberg FDR correction across time points are reported in the Supplementary Information (Supplementary Table 1); these complement the cluster-level inference by identifying individual time points at which the dose difference survived correction. To assess sensitivity to the temporal granularity of TET ratings, all LME models and cluster-based permutation tests were additionally estimated after downsampling the TET time series to non-overlapping 20 s and 30 s bins by averaging within each window.

In all time-course figures, the group mean ± s.e.m. is accompanied by the trajectory of every participant, drawn as a thin line in the modality colour for the High dose and as a thin grey dashed line for the Low dose. For the five individual affective items of Fig. 4a, whose panels are small, the spread across participants is shown as the 10th–90th percentile band for each dose instead of individual lines. Panels that display model coefficients show the fixed-effect estimate and its 95% confidence interval. The numerical values underlying every figure panel are provided as Supplementary Data 1, with one worksheet per panel and an index sheet mapping each panel to the file of the analysis pipeline that produced it.

### Physiological and affective data integration

To examine the specific relationships between physiological activation and subjective affective experience, we computed Pearson correlations between individual physiological measures (HR, SMNA AUC, and RVT) and TET-derived affective dimensions, separately for RS and DMT states. Prior to computing correlations, TET time series were averaged within the same non-overlapping 30-s windows used to summarise physiological measures (0–9 min), yielding 18 matched data points per participant per state and dose (each bin aggregating ~7.5 native 4-s samples). Univariate and composite coupling analyses were computed on the intersection subsample with complete concurrent recordings across the three modalities (*n* = 7: S04, S06, S07, S16, S18–S20; see Supplementary Fig. 6). Individual-modality correlations are reported alongside the composite Physiological Arousal Index to clarify the relative contribution of each channel while holding the subsample constant, rather than as a sample-size optimisation. Multiple comparison correction was applied across the set of physiological measures and states within each hypothesis family using the Benjamini–Hochberg FDR procedure. Three families were defined, each grouping correlations that test a related hypothesis: (1) correlations between physiological measures and the Affective Arousal Index (Emotional Intensity; 3 physiological measures × 2 states = 6 tests), (2) correlations between physiological measures and the Affective Valence Index (6 tests), and (3) correlations between physiological measures and all individual affective dimensions (6 dimensions 3 measures 2 states = 36 tests). Within each family, *p* values were jointly corrected across the constituent tests. Note that Emotional Intensity appears in both family (1), as the Affective Arousal Index proxy, and family (3), as one of the six affective dimensions; these families address related but distinct hypotheses and were corrected independently. Subsequently, to quantify the proportion of variance in subjective experience explained by the composite Physiological Arousal Index (PC1), we fit ordinary least squares (OLS) regression models predicting each TET dimension fro*_√_*mPC1, separately by state. The Pearson correlation between PC1 and each TET dimension is given by 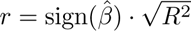.

Finally, to identify shared latent dimensions between physiological signals and subjective affective experience, we conducted Canonical Correlation Analysis (CCA). To address the repeated-measures structure (*n* = 7 participants), we implemented exact participant-level permutation testing by exhaustively enumerating all possible derangements, i.e., permutations in which no participant is paired with their own data (*D*(7) = 1,854 unique derangements for *n* = 7 participants)^93^. For each derangement, participant pairings between physiological and affective data were shuffled while preserving within-participant temporal structure. Because all *D*(7) = 1,854 derangements were evaluated, the resulting *p* values are exact, computed as (*b* + 1)*/*(*m* + 1), where *b* is the number of derangements whose canonical correlation equals or exceeds the observed value and *m* = 1,854. Canonical loadings *r >* .30 were considered substantive. Complementarily, we assessed generalisability via Leave-One-Subject-Out Cross-Validation, quantifying predictive performance as the out-of-sample canonical correlation (*r*_oos_), with significance evaluated using a one-sample *t*-test (one-tailed, *H*_1_: *r >* 0) on Fisher *Z*-transformed *r*_oos_ values across folds.

## Acknowledgements

We thank the participants who took part in this study and the facilitator who assisted them during the sessions.

## Author contributions

**T.D.:** Conceptualization, Methodology, Software, Validation, Formal analysis, Investigation, Data curation, Visualization, Project administration, Writing — Original draft, Writing — Review & editing. **T.G.G.:** Software, Formal analysis, Writing — Original draft. **J.R.C.:** Software, Formal analysis, Writing — Review & editing. **E.L.-H.:** Methodology, Investigation, Project administration, Writing — Review & editing. **C.P.:** Investigation, Project administration. **F.C.:** Investigation. **N.B.:** Investigation. **L.D.L.F.:** Investigation. **S.M.:** Investigation. **D.C.:** Investigation. **T.B.:** Conceptualization, Methodology. **D.V.:** Conceptualization, Supervision, Writing — Original draft, Writing — Review & editing. **E.T.:** Conceptualization, Methodology, Investigation, Resources, Supervision, Project administration, Funding acquisition, Writing — Original draft, Writing — Review & editing.

## Data availability

The raw multimodal physiological recordings (BrainVision format, ECG/EDA/respiration/EEG channels), the time-resolved phenomenological reports (Temporal Experience Tracing), and the preprocessed per-participant derivatives that support the findings of this study are openly available on Zenodo at https://doi.org/10.5281/zenodo.23111171. The numerical source data underlying every figure panel are provided as Supplementary Data 1 (one worksheet per panel, with an index sheet) and, as CSV files, in the code repository and in the data deposit. Per-participant age, which enters the intrinsic heart rate comparison, is withheld from the public deposit and from Supplementary Data 1 as a quasi-identifier in a sample of this size; the corresponding worksheet reports each participant’s peak heart rate and whether it exceeded the estimated intrinsic rate and the upper 95% limit of that estimate, but not the age or the estimated intrinsic rate, from which age could be recovered.

## Code availability

The analysis code that reproduces all main and supplementary figures and statistical results is available on GitHub at https://github.com/tomdamelio/dmt-emotions, with a citable archive of the version used in this paper deposited on Zenodo at https://doi.org/10.5281/zenodo.23111076. The repository also contains the script that assembles Supplementary Data 1 from the outputs of the analysis pipeline.

## Use of large language models

During the preparation of this manuscript, the authors used Claude (Anthropic) for language polishing, consistency review (e.g., British/American English harmonisation, statistical notation consistency, cross-reference verification), and preparation of the Zenodo deposit metadata. All output was verified by the authors, who take full responsibility for the content.

