## Supplementary Information for "Multimodal autonomic arousal tracks dose-dependent affective dynamics during the acute effects of DMT"

931 # Supplementary Information

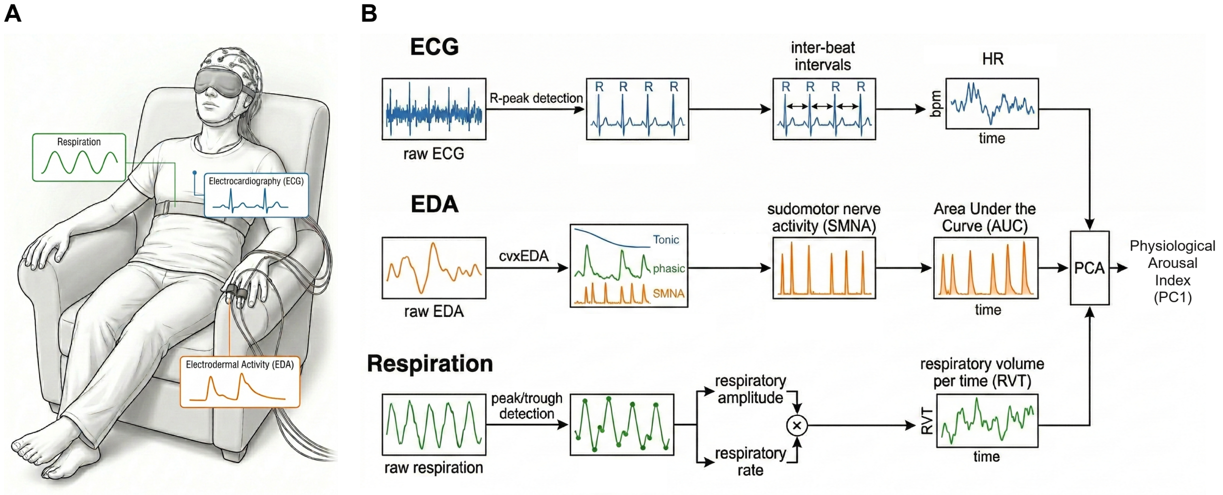


Supplementary Fig. 1: Recording setup and signal processing pipeline. **a**, Schematic of the recording setup. Participants rested in a reclined position with eyes closed under an eye mask throughout the session. Peripheral physiological signals were recorded continuously using a Brain Products amplifier (250 Hz): electrocardiography (ECG; modified lead II bipolar montage), respiration (thoracic effort belt), and electrodermal activity (EDA; finger electrodes), acquired simultaneously with EEG during both Resting State and DMT blocks. **b**, Signal processing and feature extraction. ECG, EDA, and respiration signals were processed to extract heart rate (HR), sudomotor nerve activity (SMNA), and respiratory volume per time (RVT), which were combined via principal component analysis (PCA) into the Physiological Arousal Index.

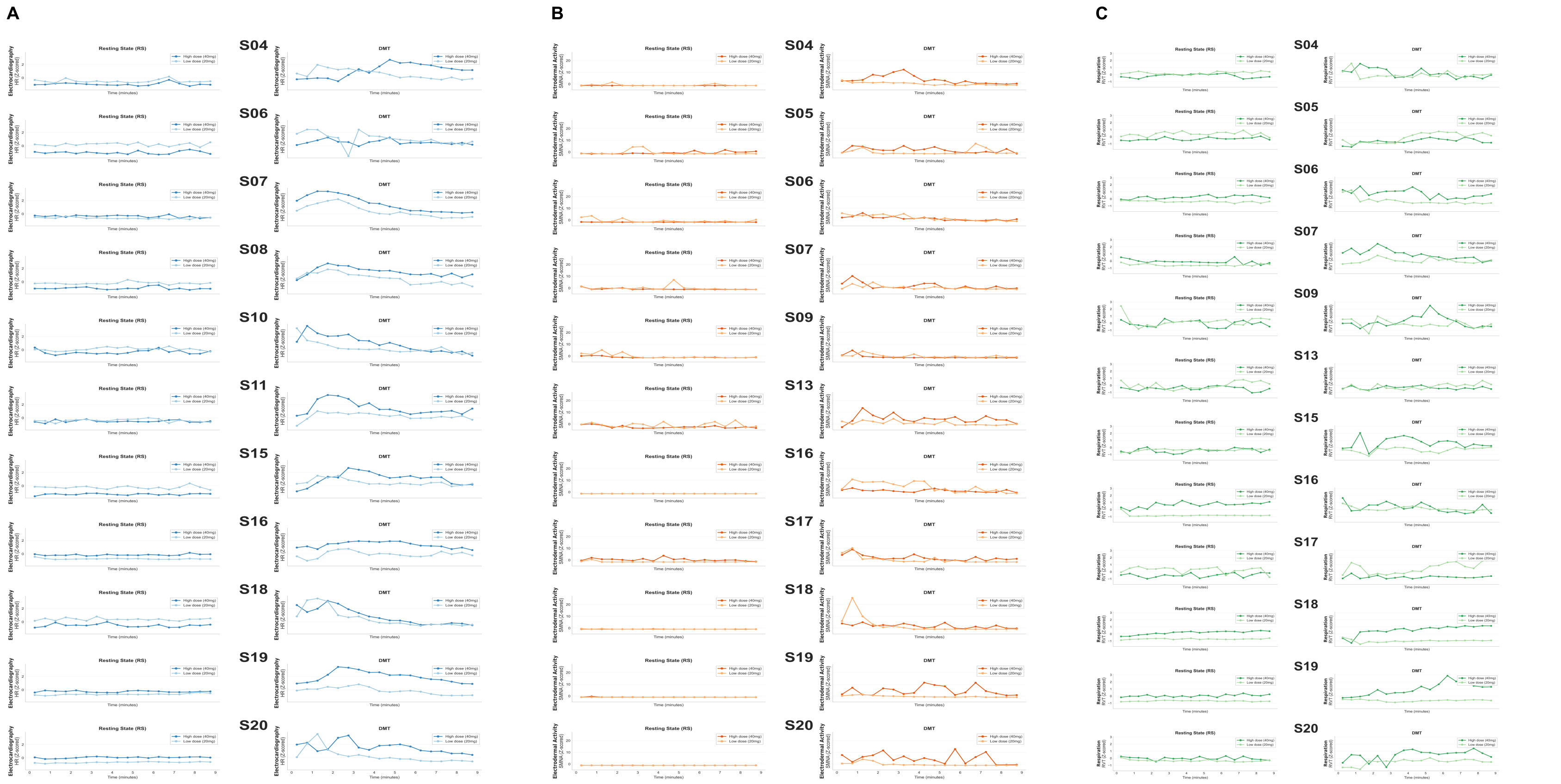


Supplementary Fig. 2: Participant-level physiological trajectories. **a–c**, Individual time courses ($z$-scored within participant; 30-s bins, 0–9 min) for Heart Rate (**a**; HR), Sudomotor Nerve Activity (**b**; SMNA), and Respiratory Volume per Time (**c**; RVT). Plots display valid recordings for each participant across Resting State (RS; left) and DMT (right) sessions, comparing Low and High doses aligned to inhalation onset ($t_0$). Included participants vary by modality due to signal quality (ECG, $n = 11$: S04, S06–S08, S10, S11, S15, S16, S18–S20; EDA, $n = 11$: S04–S07, S09, S13, S16–S20; Respiration, $n = 12$: S04–S07, S09, S13, S15–S20). Source data for these traces are the continuous per-participant signals in the derivatives folder of the data deposit (see Data availability).

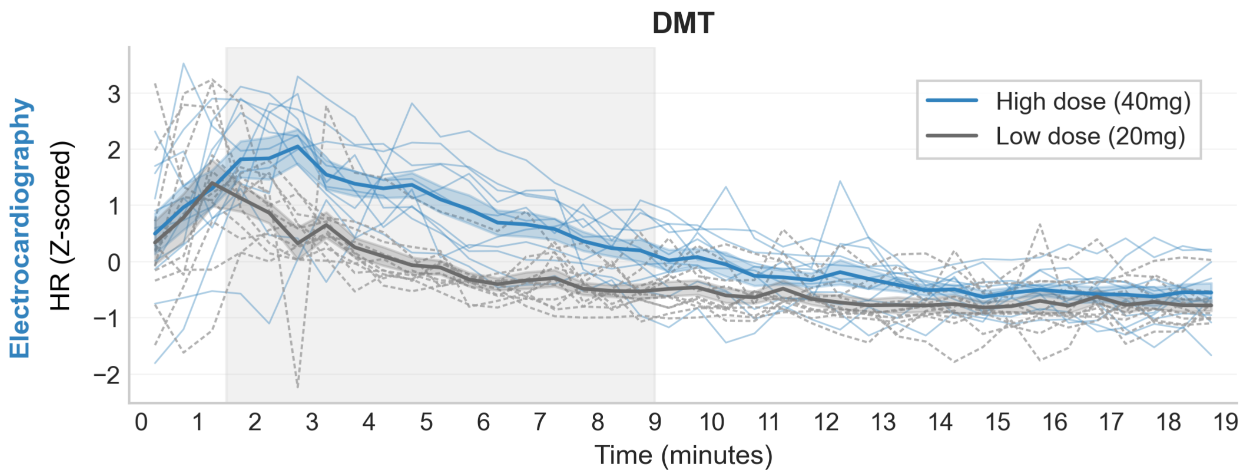


Supplementary Fig. 3: Extended heart rate trajectories. Group-averaged Heart Rate (HR; $z$-scored) time courses (mean ± s.e.m.; $n = 11$) during DMT, aligned to inhalation onset ($t_0$). The plot displays the full 0–19 min recording session, extending the restricted 0–9 min window used for the main multivariate analyses. Thin lines show each participant (High dose in colour, Low dose in grey, dashed). Shaded vertical bands indicate clusters of significant High vs. Low differences (cluster-based sign-flip permutation test, one-tailed, computed over the 38 extended windows; see Methods). The single cluster identified here (approximately 1.5–9.0 min, exact $p = .007$) coincides with the 1.5–9 min cluster of Fig. 2a in both onset and offset: extending the analysis to the full recording does not lengthen the interval over which the doses separate. Source data are provided as a Source Data file (Supplementary Data 1).

### Time-domain heart rate variability

Time-domain variability measures are reported here rather than in the main text because, as set out below, they support no claim beyond those the main text already makes on firmer ground.

RMSSD, SDNN, and RMSSD divided by the mean inter-beat interval were computed in 120-s windows sliding in 30-s steps, on Kubios-corrected R-peaks with the artefact handling described in Methods, in the ten participants with valid ECG whose four recordings all had no more than 10% invalid intervals (see Methods). Fixed effects were estimated with the same linear mixed-effects specification used throughout, carrying the by-participant random slopes described in Methods, and separation over time, both between states and between doses, was assessed with the same exact sign-flip permutation cluster test.

RMSSD was lower during DMT than at rest ($\beta = -0.42$, 95% CI $[-0.67, -0.16]$, $p = .001$) and showed a dose-dependent decrease (State × Dose: $\beta = -0.97$, 95% CI $[-1.69, -0.24]$, $p = .009$); the cluster test located the departure from rest between 1.0 and 3.0 min after inhalation (exact $p = .033$). In absolute units, median RMSSD under the High dose fell from 32.5 ms in the first window to 21.5 ms at 3.5 min, against a resting median that ranged from 44.7 to 54.7 ms across the corresponding windows, and recovered to 43.4 ms by 8 min while heart rate remained elevated. SDNN showed the same dose-dependent pattern (State × Dose: $\beta = -0.91$, 95% CI $[-1.55, -0.27]$, $p = .005$), although its State effect was not significant ($\beta = 0.29$, 95% CI $[-0.05, 0.62]$, $p = .093$). The cluster test comparing the two doses within DMT did not reach significance for either measure (RMSSD, $p = .091$ and $p = .119$ for the two candidate clusters; SDNN, $p = .069$ and $p = .245$), so the dose dependence is supported by the model over the 9-min window but not by any specific contiguous epoch. Heart rate in this subsample did separate the doses, with a cluster at approximately 2.5–8 min (exact $p = .006$, $n = 10$; Supplementary Fig. 4d).

The interpretation, however, is limited by the dependence of RMSSD on cycle length. Within participants, RMSSD and the mean inter-beat interval were strongly correlated across DMT windows (median $r = .81$, interquartile range .55–.92), and the log–log slope of RMSSD on mean interval had a median of 2.16, so absolute variability fell more than proportionally as the cycle shortened. When RMSSD was expressed relative to the mean inter-beat interval, neither the State effect ($\beta = 0.01$, $p = .969$) nor the State × Dose interaction ($\beta = -0.68$, $p = .100$) was detectable, and no cluster survived. Dividing by the mean interval is a conservative correction, since vagal withdrawal itself shortens the interval and the ratio therefore discounts some genuinely vagal variation. Absolute and normalised RMSSD are reported together because the present data cannot separate the vagal contribution to variability from the contribution of cycle length. We also found no dose-dependent late vagal rebound: between 10 and 19 min the High-minus-Low difference in RMSSD was 0.30 in $z$ units with 6 of 10 participants positive ($p = .434$).

Because of these limitations, the main text rests its claim about sympathetic engagement on the intrinsic-rate comparison, which is computed from heart rate in absolute units and does not depend on any variability measure.

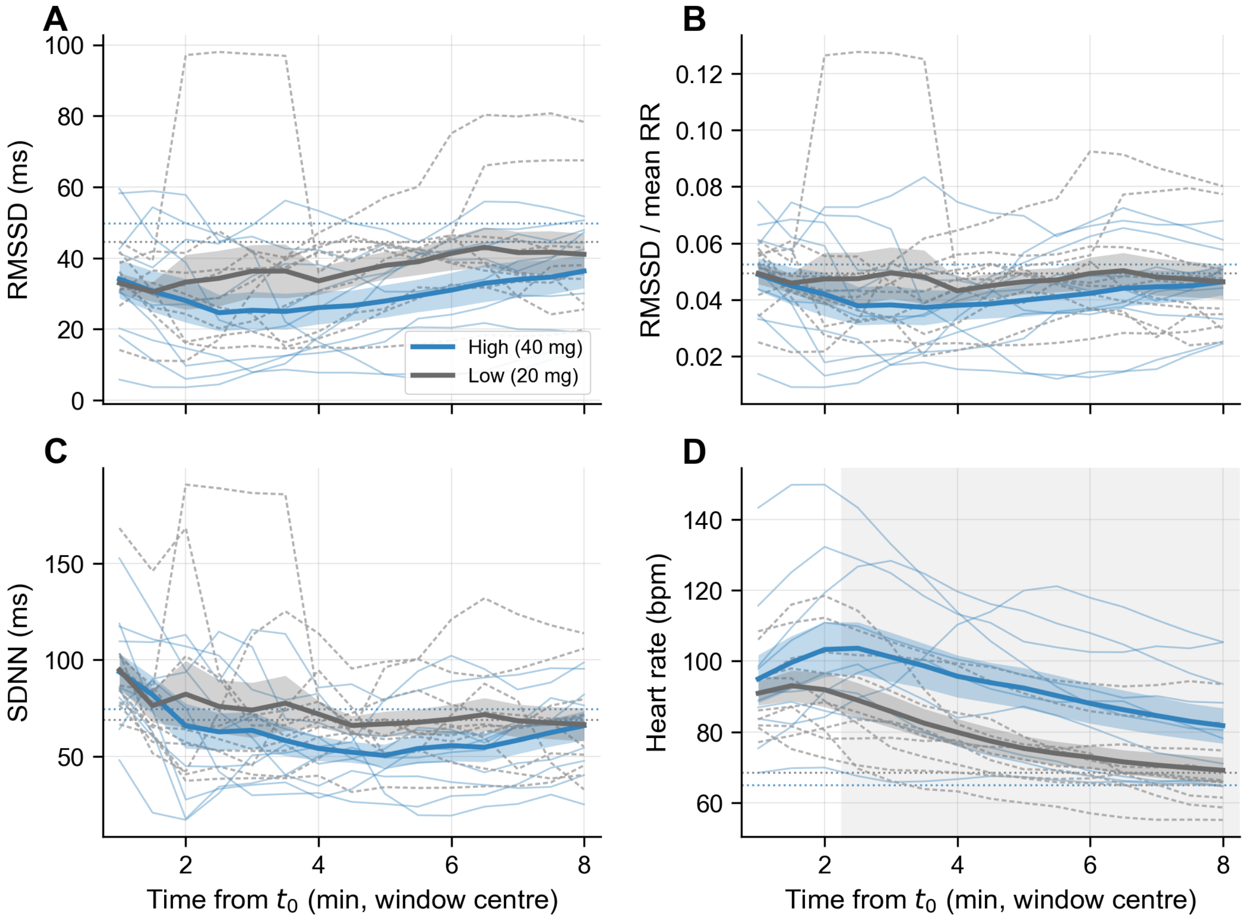


Supplementary Fig. 4: Time-domain heart rate variability during DMT. **a**, RMSSD; **b**, RMSSD divided by the mean inter-beat interval; **c**, SDNN; **d**, heart rate in beats per minute. All measures were computed in 120-s windows sliding in 30-s steps and are plotted at the window centre. Thick lines and shading give the group mean ± s.e.m. ($n = 10$); thin lines show each participant (High dose in colour, Low dose in grey, dashed). Dotted horizontal lines give the resting-state mean of the corresponding dose. The shaded rectangle in **d** indicates the one cluster of significant High vs. Low separation (exact sign-flip permutation cluster tests, one-tailed; see Methods), and the variability measures in **a–c** show no dose cluster. The decrease of RMSSD below its resting level reported in the text (1.0–3.0 min) comes from a DMT vs. RS comparison with the two doses averaged and is therefore not shaded. Source data are provided as a Source Data file (Supplementary Data 1).

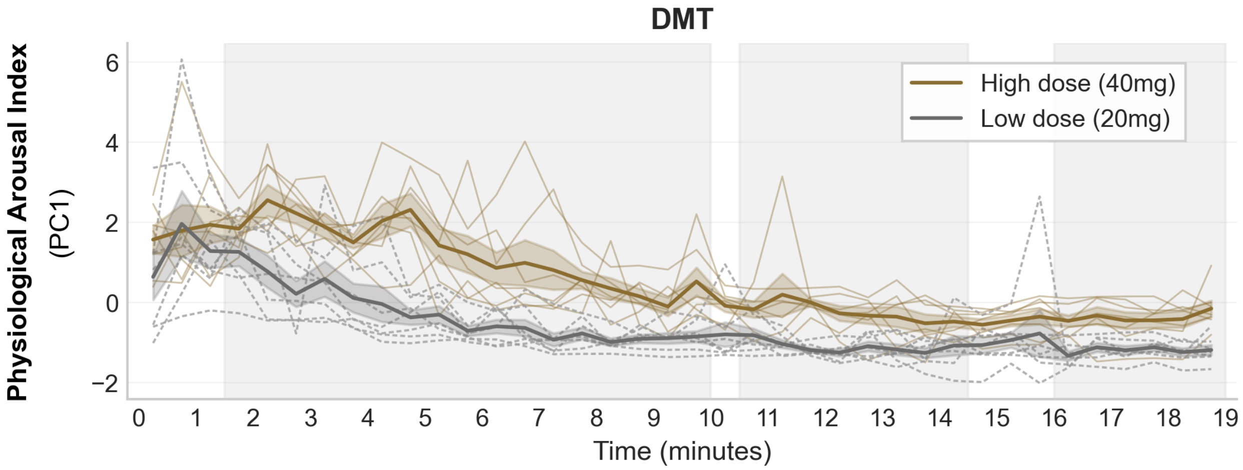


Supplementary Fig. 5: Extended Physiological Arousal Index trajectories. Group-averaged time courses (mean $\pm$ s.e.m.; $n = 7$) of the Physiological Arousal Index ($z$-scored PC1) during DMT. Data are aligned to inhalation onset ($t_0$) and displayed over the full 0–19 min window to extend the primary analysis. Thin lines show each participant (High dose in colour, Low dose in grey, dashed). Shaded vertical bands indicate clusters of significant High vs. Low differences (cluster-based sign-flip permutation test, one-tailed, computed over the 38 extended windows; exact $p = .023$ for each of the three clusters; see Methods). Source data are provided as a Source Data file (Supplementary Data 1).

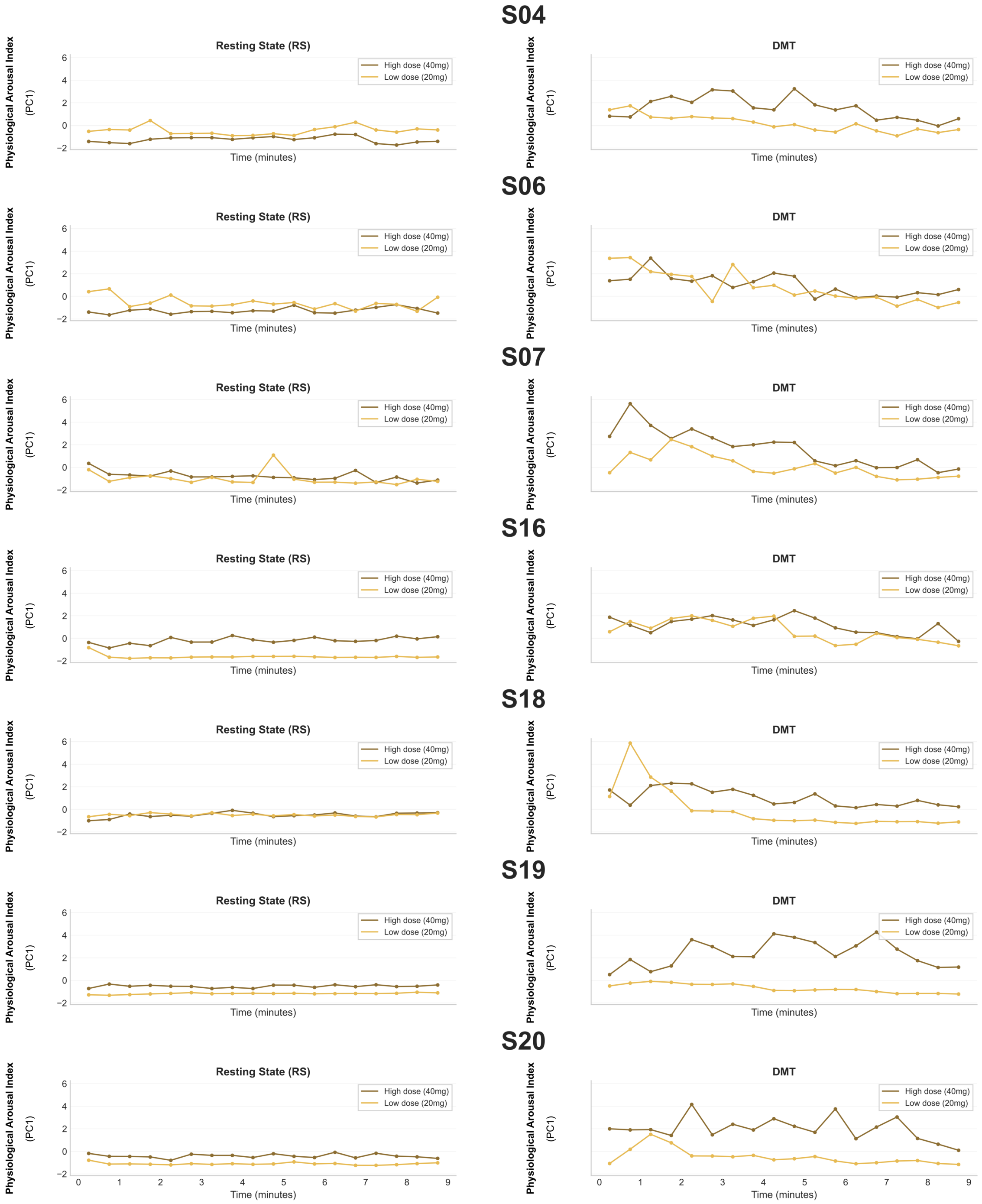


Supplementary Fig. 6: Participant-level trajectories of the Physiological Arousal Index. Individual time courses (30-s bins) of the Physiological Arousal Index ($z$-scored PC1) for the subset of participants with complete concurrent physiological recordings ($n = 7$: S04, S06, S07, S16, S18–S20). Panels display Resting State (RS) and DMT sessions for Low and High doses, aligned to inhalation onset ($t_0$). The index represents the first principal component capturing shared variance across HR, SMNA, and RVT. Source data are provided as a Source Data file (Supplementary Data 1).

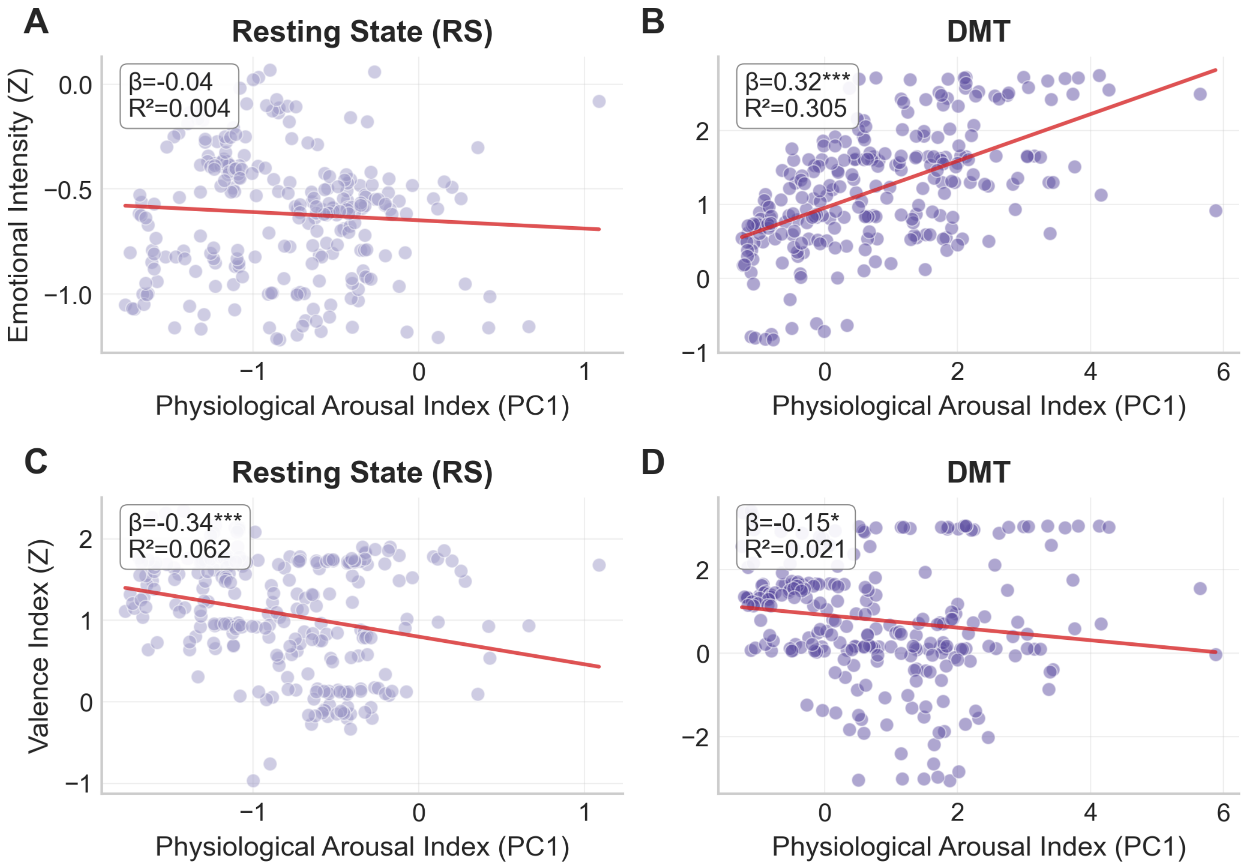


Supplementary Fig. 7: State-dependent associations between physiological arousal and affective dimensions. Scatterplots displaying the relationship between the Physiological Arousal Index (PC1; x-axis) and affective ratings (y-axis). Each point represents a single 30-s bin; red lines indicate ordinary least-squares fits; asterisks on the in-panel coefficients denote * $p < .05$ and *** $p < .001$. **a, b**, Emotional Intensity ($z$-scored) during Resting State (**a**) and DMT (**b**; $\beta = 0.32$, $R^2 = .30$, $p < .001$). **c, d**, Affective Valence Index (Pleasantness minus Unpleasantness; $z$-scored) during Resting State (**c**; $\beta = -0.34$, $p < .001$) and DMT (**d**; $\beta = -0.15$, $p = .021$). Note that this regression on the composite Physiological Arousal Index differs from the per-modality correlations reported in the main text, where HR was the only individual modality showing a significant association with valence during DMT. Source data are provided as a Source Data file (Supplementary Data 1).

Supplementary Table 1: Cluster-based permutation results and window-wise FDR-corrected comparisons for all shaded time courses. Cluster extents are approximate (see Methods). Cluster-level $p$ values are exact, obtained by exhaustive sign-flip permutation ($2^n$ permutations for one-tailed tests and $2^{n-1}$ for two-tailed ones). FDR windows list the time points at which the paired $t$-test survived Benjamini–Hochberg correction across time points ($p_{\mathrm{FDR}} < .05$). Physiological time courses use 30-s windows; TET time courses use native 4-s samples (DMT: 0–20 min; RS: 0–10 min). Resting-state comparisons are listed even when neither test returns a significant result, because the manuscript reports the absence of dose separation at rest.

| Measure | Comparison | Tail | $n$ | Cluster(s), min | FDR windows, min |
|---|---|---|---|---|---|
| Heart rate | DMT | one-tailed | 11 | 1.5–9.0 ($p = .0015$) | 1.5–9.0 |
| Heart rate | RS | two-tailed | 11 | none | none |
| Heart rate | DMT, 0–19 min | one-tailed | 11 | 1.5–9.0 ($p = .007$) | 2.5–3.0; 3.5–9.0 |
| SMNA | DMT | one-tailed | 11 | 4.0–5.5 ($p = .041$) | none |
| SMNA | RS | two-tailed | 11 | none | none |
| RVT | DMT | one-tailed | 12 | 1.0–5.0 ($p = .011$) | 1.5–5.0 |
| RVT | RS | two-tailed | 12 | none | none |
| Physiological Arousal Index | DMT | one-tailed | 7 | 2.0–9.0 ($p = .016$) | 2.0–3.0; 3.5–9.0 |
| Physiological Arousal Index | RS | two-tailed | 7 | none | none |
| Physiological Arousal Index | DMT, 0–19 min | one-tailed | 7 | 1.5–10.0 ($p = .023$); 10.5–14.5 ($p = .023$); 16.0–19.0 ($p = .023$) | 2.0–3.0; 3.5–10.0; 10.5–14.5; 16.0–19.0 |
| TET: Emotional Intensity | DMT High vs. Low | two-tailed | 18 | 0.3–10.1 ($p = .025$) | 1.1–3.2 |
| TET: Emotional Intensity | DMT vs. RS | two-tailed | 18 | 0.0–9.9 ($p < .001$) | 0.0–9.9 |
| TET: Interoception | DMT vs. RS | two-tailed | 18 | 0.0–9.9 ($p < .001$) | 0.0–9.9 |
| TET: Anxiety | DMT vs. RS | two-tailed | 18 | 0.0–9.9 ($p < .001$) | 0.0–9.9 |
| TET: Unpleasantness | DMT vs. RS | two-tailed | 18 | 0.0–9.9 ($p < .001$) | 0.0–9.9 |
| TET: PC1 | DMT High vs. Low | two-tailed | 18 | 0.7–11.4 ($p = .021$) | 1.0–7.6 |
| TET: PC1 | DMT vs. RS | two-tailed | 18 | 0.0–9.9 ($p < .001$) | 0.0–9.9 |

Supplementary Table 2: Sensitivity of LME State × Dose estimates to TET temporal resolution.

| Outcome | 4 s (original) | 20 s | 30 s |
|---|---|---|---|
| Arousal (Emotional Intensity) | $0.49\ [-0.01,\ 0.99]^{\mathrm{ns}}$ | $0.49\ [-0.01,\ 0.99]^{\mathrm{ns}}$ | $0.49\ [-0.01,\ 0.99]^{\mathrm{ns}}$ |
| Valence (Pleasantness − Unpleasantness) | $-0.81\ [-1.50,\ -0.12]^{*}$ | $-0.81\ [-1.51,\ -0.11]^{*}$ | $-0.81\ [-1.51,\ -0.11]^{*}$ |
| *Individual dimensions* | | | |
| Interoception | $0.50\ [0.02,\ 0.98]^{*}$ | $0.50\ [0.02,\ 0.98]^{*}$ | $0.50\ [0.02,\ 0.98]^{*}$ |
| Anxiety | $0.42\ [-0.05,\ 0.88]^{\mathrm{ns}}$ | $0.42\ [-0.05,\ 0.88]^{\mathrm{ns}}$ | $0.42\ [-0.05,\ 0.88]^{\mathrm{ns}}$ |
| Unpleasantness | $0.62\ [0.23,\ 1.01]^{**}$ | $0.62\ [0.23,\ 1.02]^{**}$ | $0.62\ [0.23,\ 1.02]^{**}$ |
| Pleasantness | $-0.19\ [-0.74,\ 0.37]^{\mathrm{ns}}$ | $-0.19\ [-0.74,\ 0.37]^{\mathrm{ns}}$ | $-0.19\ [-0.74,\ 0.37]^{\mathrm{ns}}$ |
| Bliss | $0.10\ [-0.60,\ 0.80]^{\mathrm{ns}}$ | $0.10\ [-0.60,\ 0.80]^{\mathrm{ns}}$ | $0.10\ [-0.60,\ 0.81]^{\mathrm{ns}}$ |

*Notes.* Fixed-effect coefficients ($\beta$) and 95% confidence intervals for the State × Dose interaction term (`state[T.DMT]:dose[T.High]`) from models re-estimated after downsampling TET time series to 20 s and 30 s non-overlapping bins. The 4 s column corresponds to the original TET sampling resolution used in the main analyses. Point estimates and intervals are essentially identical across resolutions (300 → 60 → 40 time points for DMT; 150 → 30 → 20 for RS), because the models carry a by-participant random intercept and slopes and the uncertainty of the interaction reflects variation between participants rather than the number of time points. Reference levels: State = RS, Dose = Low (20 mg). Significance (FDR-corrected, as in Fig. 4b): $^{**}p < .01$, $^{*}p < .05$, ns = not significant.

Supplementary Table 3: Quality assessment of individual physiological recordings.

| | DMT High | | | RS High | | | DMT Low | | | RS Low | | |
|---|---|---|---|---|---|---|---|---|---|---|---|---|
| Participant | EDA | ECG | Resp | EDA | ECG | Resp | EDA | ECG | Resp | EDA | ECG | Resp |
| S01 | M | M | M | M | M | M | G | G | A | G | G | A |
| S02 | G | A | A | G | A | A | M | M | M | M | M | M |
| S03 | G | G | A | G | G | G | M | M | M | M | M | M |
| S04 | G | G | A | G | G | A | G | A | A | G | G | G |
| S05 | A | B | G | G | G | G | A | G | A | A | G | G |
| S06 | G | A | A | A | G | A | G | A | A | A | G | A |
| S07 | G | G | A | G | G | G | A | G | A | A | G | A |
| S08 | B | G | B | B | G | B | A | A | A | G | G | G |
| S09 | G | B | A | G | G | A | G | A | G | G | G | G |
| S10 | G | G | A | G | G | A | B | G | B | B | A | G |
| S11 | B | G | B | B | G | B | G | G | A | G | A | A |
| S12 | B | B | B | B | G | B | G | G | G | A | A | G |
| S13 | A | B | A | A | B | A | G | B | A | G | B | G |
| S15 | G | G | G | A | G | G | B | G | G | B | G | G |
| S16 | G | G | G | A | G | G | G | A | G | A | A | G |
| S17 | G | G | A | G | G | G | G | G | G | G | B | G |
| S18 | G | G | G | A | A | G | G | G | G | G | G | G |
| S19 | G | G | G | G | G | G | G | G | G | G | G | G |
| S20 | G | A | G | G | G | G | G | G | G | G | G | G |

*Notes.* Quality ratings for Electrodermal Activity (EDA), Electrocardiography (ECG), and Respiration across four conditions: High dose (40 mg) and Low dose (20 mg) DMT, and their respective same-day Resting State (RS) baselines. Signals were visually inspected for signal-to-noise ratio and physiological plausibility. **G**, Good (high quality, suitable for analysis); **A**, Acceptable (usable with minor limitations); **B**, Bad (excluded due to excessive noise); **M**, Missing (data unavailable). Only participants with valid recordings (ratings G or A) were retained for the corresponding group-level analyses ($n = 19$ enrolled). Participant identifier S14 is not included; the final sample comprised 19 participants (S01–S13, S15–S20).
